# Can deep learning predict human intelligence from structural brain MRI?

**DOI:** 10.1101/2023.02.24.529924

**Authors:** Mohammad Arafat Hussain, Danielle LaMay, Ellen Grant, Yangming Ou

**Affiliations:** Department of Pediatrics, Boston Children’s Hospital, Harvard Medical School, 401 Park Drive, Boston, MA 02115, USA; Khoury College of Computer and Information Science, Northeastern University, 360 Huntington Ave, Boston, MA 02115, USA; Computational Health Informatics Program, Boston Children’s Hospital, Harvard Medical School, 401 Park Drive, Boston, MA 02115, USA; Department of Radiology, Harvard Medical School, 401 Park Drive, Boston, MA 02115, USA

## Abstract

Can brain structure predict human intelligence? T1-weighted structural brain magnetic resonance images (sMRI) have been correlated with intelligence. Nevertheless, population-level association does not fully account for individual variability in intelligence. To address this, individual prediction studies emerge recently. However, they are mostly on predicting fluid intelligence (the ability to solve new problems). Studies are lacking to predict crystallized intelligence (the ability to accumulate knowledge) or general intelligence (fluid and crystallized intelligence combined). This study tests whether deep learning of sMRI can predict an individual subject’s verbal, comprehensive, and full-scale intelligence quotients (VIQ, PIQ, FSIQ), which reflect both fluid and crystallized intelligence. We performed a comprehensive set of 432 experiments, using different input images, six deep learning models, and two outcome settings, on 850 autistic and healthy subjects 6-64 years of age. Results show promise with statistical significance, and also open up questions inviting further future studies.

## Introduction

Human intelligence is influenced by nature and nurture. The former is coded in the gene. The latter relates to the environment, nutrition, socioeconomics, lifestyle, and other factors. Nature and nurture jointly shape the human brain, which can be non-invasively observed by magnetic resonance imaging (MRI)^1^. A mystery is, can MRI explain individual variability in human intelligence?

Among MRI sequences, structural MRI (sMRI) measures neuroanatomy and is available in almost every MRI scan. sMRI metrics, such as regional volumes, cortical thickness, and cortical surface areas, are correlated with human intelligence^2–4^. These population-level studies, however, do not explain individual variability in intelligence^5^.

Recent studies have started to use the machine especially deep learning to predict intelligence for individuals but left many open questions. In over 20 studies corresponding to a 2019 Grand Challenge, the predicted fluid intelligence had a mean square error ranging from 86 to 103 (for a range of true residual fluid intelligence score of [−40, 30])^6–22^. The modest accuracy may suggest the need for more sophisticated deep learning algorithms. Or, perhaps sMRI does not contain sufficient information for fluid intelligence, the ability to solve new problems that peak in young adulthood. Can sMRI predict an individual’s crystallized intelligence, the ability to accumulate knowledge that grows with age? Can sMRI predict an individual’s general intelligence (fluid and crystallized intelligence combined)? Does the accuracy vary by deep learning algorithms? What neuroanatomy does deep learning rely on to make the prediction?

This paper aimed to address these open questions. We conducted comprehensive experiments quantifying the accuracy of predicting an individual’s verbal, performance, and full-scale intelligence quotients (VIQ, PIQ, and FSIQ), which are loosely related to crystallized, fluid, and general intelligence^23^. We used different 2D and 3D deep learning algorithms, with different input sMRI channels, different parameter settings, and different prediction modes, on sMRI from 800+ individuals 6-64 years of age. Results offer new insights into the extent sMRI can infer an individual’s intelligence.

## Results

### Experimental Setup

We trained two 2D and four 3D deep CNNs (listed in Fig. 1(b)) using T1-weighted MRI volumes (*N* = 850) in two settings. In the first setting, we used intensity (i.e., **contrast** channel) and RAVENS (regional analysis of volumes examined in normalized space; i.e., **morphometry** channel)^24^ images to predict three IQ scores separately (i.e., FSIQ or PIQ, or VIQ). On the other hand, we used intensity and RAVENS maps to predict three IQ scores simultaneously (i.e., FSIQ and PIQ, and VIQ) in the second setting (see Fig. 1(c)). For 2D CNNs, we chose a different number of axial slices (*n* = [5, 10, 20, 40, 70, 100, 130]) as channels. Since our 3D volume has 130 axial slices, we selected slices as [65 − ⌊*n/*2⌋, 65 + ⌊*n/*2⌋]. Intensity (denoted as I) and RAVENS (denoted as R) maps were used as inputs to both 2D and 3D CNNs separately and together as channels (denoted as IR). We used 5-fold cross-validation for each experiment, and each fold was scheduled to run for 30 and 100 epochs for 2D and 3D CNNs, respectively. We ran a total of 432 experiments, which involve training deep CNN training in each case. We picked the best validation performance in each fold.

**Figure 1.**
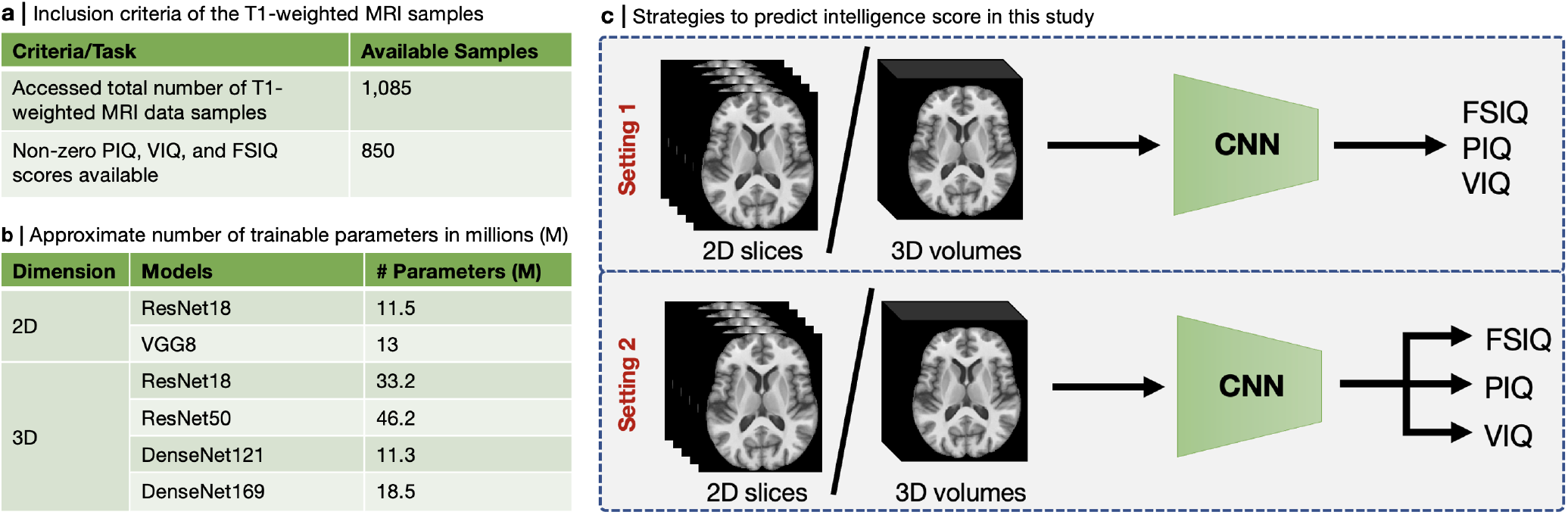
(a) Inclusion criteria of T1-weighted MRI samples in this study, (b) the approximate number of trainable parameters in millions (M) of the six deep CNNs used in this study, and (c) schematic diagram of our intelligence score prediction strategies.

### Absolute and Residual IQ Prediction by 2D CNNs

We ran 336 experiments using 2D CNNs for seven different slice numbers, three types of input, and two output settings. We presented individual experiment-wise IQ score prediction performance in terms of Pearson correlation (*r*) and mean absolute error in Supplementary Fig. 1 and Tables 1–8, where we observe that statistically significant (*p* < 0.001) PIQ, VIQ, and FSIQ prediction performance are achieved by 2D-ResNet18 for contrast input (i.e., I) and absolute IQ scores in setting 1. Nonetheless, we present an overall absolute and residual IQ prediction performance in terms of *r* for different slice numbers and input types in Fig. 2(a, b) and (c, d), respectively. We see in Fig. 2(a, b) that the mean *r* between the ground truth and predicted absolute and residual PIQ, VIQ, and FSIQ tends to get overall better for higher slice numbers (i.e., 70, 100, and 130). Furthermore, we see in Fig. 2(c, d) that the mean *r* between the ground truth and predicted absolute and residual PIQ, VIQ, and FSIQ is the overall best for contrast input (i.e., I).

**Table 1.** Comparative best PIQ, VIQ, and FSIQ prediction performance in terms of Pearson correlation (*r*) for 2D and 3D CNNs. Associated experiment numbers (Exp. No.), mean absolute error (MAE), and mean absolute percentage error (MAPE) are also shown. * indicates a statistical significance for *p* < 0.001.

| CNN | Exp. No.<br>(PIQ/VIQ/FSIQ) | PIQ |  |  | VIQ |  |  | FSIQ |  |  |
| --- | --- | --- | --- | --- | --- | --- | --- | --- | --- | --- |
| | | $r$ | MAE | MAPE (%) | $r$ | MAE | MAPE (%) | $r$ | MAE | MAPE (%) |
| 2D | 327/24/132 | 0.232 | 10.57±9.43 | 9.58±3.16 | 0.187* | 11.82±9.19 | 11.42±10.66 | 0.216* | 11.37±8.79 | 10.85±9.38 |
| 3D | 338/424/347 | 0.191* | 11.79±9.95 | 11.77±12.71 | 0.168* | 11.54±9.50 | 98.34±49.58 | 0.192* | 11.67±9.16 | 11.40±11.25 |

**Figure 2.**
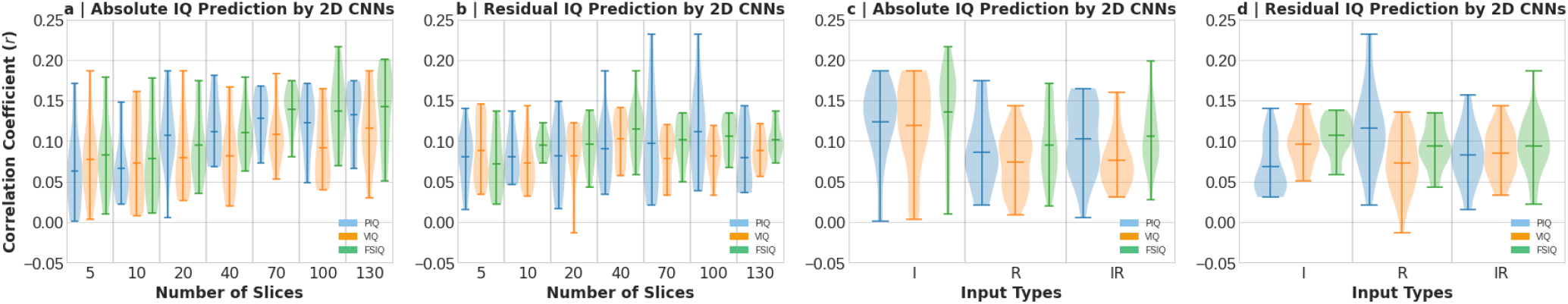
Violin plots showing absolute and residual IQ prediction performance in terms of Pearson correlation coefficient (*r*) by both 2D-ResNet18 and 2D-VGG8 in settings 1 and 2. Correlation between the ground truth and predicted (a) absolute PIQ, VIQ, and FSIQ scores *vs*. numbers of slices, (b) residual PIQ, VIQ, and FSIQ scores *vs*. numbers of slices, (c) absolute PIQ, VIQ, and FSIQ scores *vs*. input types, and (d) residual PIQ, VIQ, and FSIQ scores *vs*. input types.

### Absolute and Residual IQ Prediction by 3D CNNs

We also performed 96 experiments using 3D CNNs for three types of input and two output settings. We presented individual experiment-wise IQ score prediction performance in terms of Pearson correlation (*r*) and mean absolute error in Supplementary Fig. 2 and Tables 9–24, where we observe that better *r* with statistically significant (*p* < 0.001) for PIQ, VIQ and FSIQ scores is achieved by 3D-ResNet18 for contrast input (i.e., I). We also present an overall absolute and residual IQ prediction performance by 3D CNNs in terms of *r* for different input types in Fig. 3(a). The figure shows that the mean *r* between the ground truth and the predicted absolute PIQ, VIQ, and FSIQ is the best overall for the input of morphometry (i.e., R). Furthermore, we see in Fig. 3(a) that the mean *r* between the ground truth and predicted residual PIQ, VIQ, and FSIQ is the overall best for the contrast input (i.e., I).

**Figure 3.**
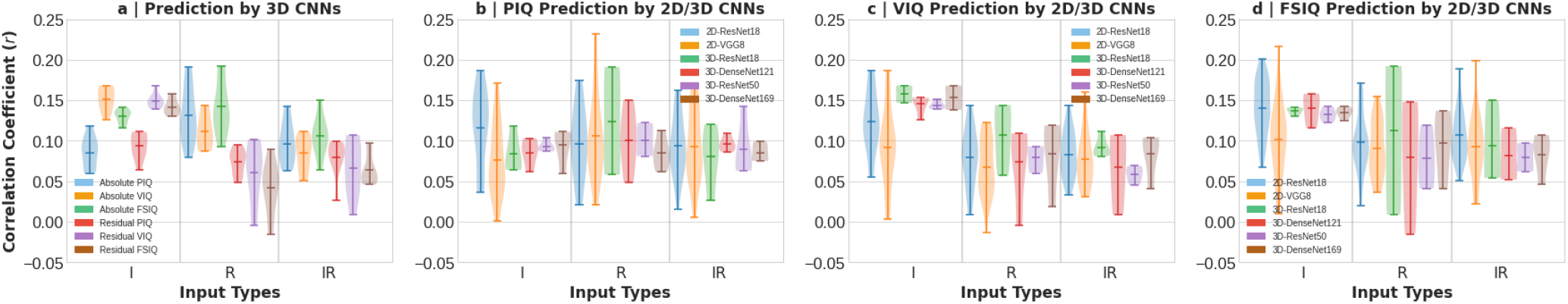
Violin plots showing absolute and residual IQ prediction performance in terms of Pearson correlation coefficient (*r*) by 2D and 3D CNNs in settings 1 and 2. (a) Correlation between the ground truth and absolute/residual PIQ, VIQ, and FSIQ scores *vs*. input types by 3D-ResNet18, 3D-ResNet50, 3D-DenseNet121, and 3D-DenseNet169. (b-d) Correlation between the ground truth and actual/residual PIQ, VIQ, and FSIQ scores *vs*. input types, respectively, by different 2D and 3D CNNs.

### Comparative IQ Prediction Performance by 2D and 3D CNNs

We present comparative PIQ, VIQ, and FSIQ prediction performance in terms of Pearson correlation (*r*) in Fig. 3(b), (c), and (d), respectively, by different 2D and 3D CNNs. In Fig. 3(b), we see that overall best PIQ prediction is achieved by 2D and 3D CNNs for the morphometry (i.e., R), followed by the contrast (i.e., I) and contrast-morphometry combined (i.e., IR) inputs. Furthermore, we see in Fig. 3(c) that overall best VIQ prediction is achieved by 2D and 3D CNNs for the contrast (i.e., I), followed by the morphometry (i.e., R) and the contrast-morphometry combined (i.e., IR) inputs. Similarly, we see in Fig. 3(d) that overall best FSIQ prediction is achieved by 2D and 3D CNNs for the contrast (i.e., I), followed by the morphometry (i.e., R) and the contrast-morphometry combined (i.e., IR) inputs. We also show a comparison of the best PIQ, VIQ, and FSIQ predictions in terms of Pearson correlation (*r*) for 2D and 3D CNNs in Table 2. We also present the associated experiment numbers (see Supplementary Tables 1–24 for cross-referencing), mean absolute error (MAE), and mean absolute percentage error (MAPE). We see in this table that the 2D CNN outperformed the 3D CNN in terms of *r*. Similarly, the 2D CNN outperformed the 3D CNN in terms of MAE and MAPE metrics, except for the MAE of VIQ score.

**Table 2.** Comparative best PIQ, VIQ, and FSIQ prediction performance in terms of Pearson correlation (*r*) for 2D and 3D CNNs. Associated experiment numbers (Exp. No.), mean absolute error (MAE), and mean absolute percentage error (MAPE) are also shown. * indicates a statistical significance for *p* < 0.001.

| CNN | PIQ |  |  | VIQ |  |  | FSIQ |  |  |
| --- | --- | --- | --- | --- | --- | --- | --- | --- | --- |
| | $r$ | MAE | MAPE (%) | $r$ | MAE | MAPE (%) | $r$ | MAE | MAPE (%) |
| 2D Resnet18 | 0.232 | 10.57±9.43 | 9.58±3.16 | 0.187* | 11.82±9.19 | 11.42±10.66 | 0.216* | 11.37±8.79 | 10.85±9.38 |
| 3D ResNet18 | 0.191* | 11.79±9.95 | 11.77±12.71 | 0.168* | 11.54±9.50 | 98.34±49.58 | 0.192* | 11.67±9.16 | 11.40±11.25 |

### Interpretation

Neuroimaging studies proposed several theories on the mapping between brain structure and function that underlie human intelligence. For example, the Parieto-Frontal Integration Theory (P-FIT)^25^ is a popular one, which assumes that humans first collect and process sensory information predominantly in the *occipital* and *temporal* areas. In the next stage, structural symbolism, abstraction, and elaboration of the basic sensory information happen in the *angular gyrus, supramarginal gyrus*, and *superior parietal* lobule. The next stage involves the interaction between *parietal areas* and *frontal lobes*. This interaction supports problem-solving, evaluation, and hypothesis testing. Once the best solution is reached, the *anterior cingulate* gets engaged in the final stage for response selection and inhibition of competing responses.

In this study, we also expect that our deep CNNs focus on P-FIT-described prominent brain regions in predicting IQ scores. We used gradient-based class activation mapping (GradCAM)^26^ to observe the most salient areas in input images/volumes, which 2D/3D CNNs have used to predict IQ scores. Since we used six deep CNNs in this study, we randomly chose a single or a few subjects per best IQ score predicting experiment per CNN. We chose experiments 49, 132, 338, 352, 364, and 376 for 2D-ResNet18, 2D-VGG8, 3D-ResNet18, 3D-ResNet50, 3D-DenseNet121, and 3D-DenseNet169, respectively.

#### Interpretation for 2D CNNs

In Fig. 4(a), we show several GradCAM-based saliency maps on axial slices for 2D-ResNet18 and 2D-VGG8 models. We see that 2D-ResNet18 focused on the part of the frontal and parietal lobes in predicting the absolute FSIQ scores for the four subjects (that is, subjects 1-4). Furthermore, we see that 2D-VGG8 focused on the occipital lobe (for subjects 5, 6, and 8) and frontal lobe (for subjects 6 and 7) to predict absolute FSIQ scores. Thus, the most salient brain regions picked up by our 2D CNNs in this study correspond to several prominent brain regions described by the P-FIT model.

**Figure 4.**
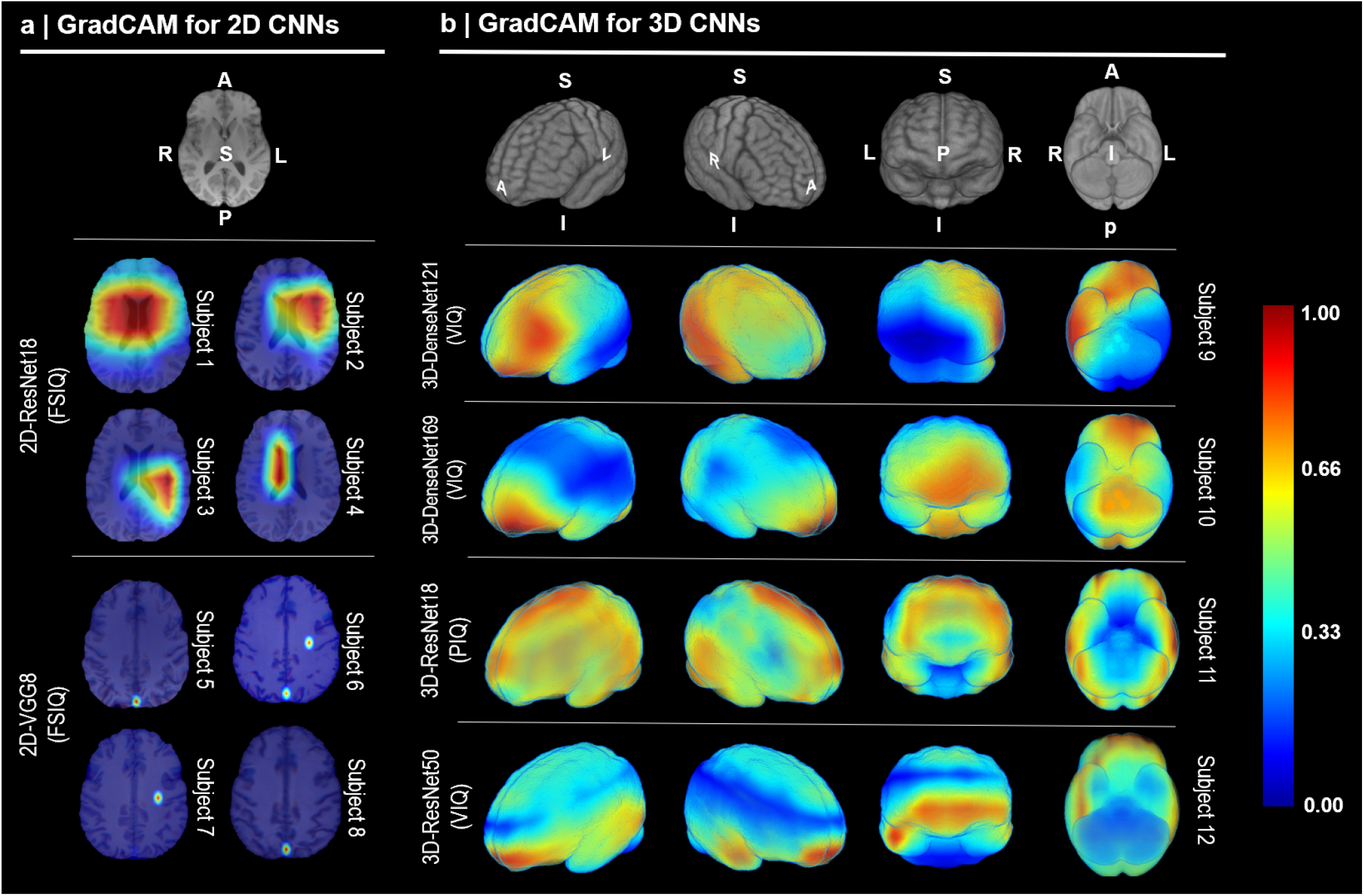
Interpretation of (a) 2D CNN-based and (b) 3D CNN-based IQ predictions using GradCAM. Acronyms– L: left, R: right, S: superior, I: inferior, A: anterior, and P: posterior.

#### Interpretation for 3D CNNs

We also show several GradCAM-based volume-rendered saliency maps for 3D-DenseNet121, 3D-DenseNet169, 3D-ResNet18, and 3D-ResNet50 in Fig. 4(b). We see in Fig. 4(b) that 3D-DenseNet121 focused on the left frontal and right parietal and temporal lobes to predict the absolute VIQ scores for subject 9. Similarly, we see in the figure that the 3D-DenseNet169 prominently focused on the frontal and occipital lobes as well as the cerebellum in predicting the absolute VIQ scores for subject 10. Furthermore, we see in Fig. 4(b) that the 3D-ResNet18 focused on almost the entire frontal, parietal, temporal, and occipital lobes as well as a part of cerebellum in predicting the absolute PIQ score for subject 11. Finally, we see in the figure that the 3D-ResNet50 focused on the frontal and parietal lobes in predicting the absolute VIQ scores for subject 12. Thus, here also, the most salient brain regions picked up by our 3D CNNs correspond to several prominent brain regions described by the P-FIT model.

## Discussion

In this study, the Pearson correlation coefficient (*r*) seems more reliable than the mean absolute error metric. The distribution of FSIQ, PIQ, and VIQ scores in our dataset follow a Gaussian-like distribution. As a result, a central tendency of the predicted scores towards the mean results in a low mean absolute error, although predictions were often inaccurate. Therefore, we considered *r* as a better indicator of the accuracy of the predicted IQ scores. In addition, the associated *p* value indicates the statistical significance of the estimated correlation.

We used three input types such as intensity (I), RAVENS (R), and a combination of I and R (IR) in our 2D and 3D CNN-based experiments. An intensity map contains contrast and textural information about the brain. On the other hand, a RAVENS map contains morphometric information that indicates the degree of volumetric deformation required for that particular brain to be registered to the brain atlas. Our results demonstrated that IQ prediction performance is overall best for the contrast input type (i.e., I) for both 2D and 3D CNNs as evident from Figs. 2(c, d), and 3.

We also used seven different slice numbers in our 2D CNN-based experiments. Our results in Fig. 2(a, b) depicted that overall absolute and residual IQ prediction performance in terms of correlation coefficient (*r*) gets improved with the increase of the number of axial slices in 2D CNNs.

In Fig. 3(b-d), we showed violin plots of different CNN-wise absolute and residual IQ prediction performance in terms of *r* for input types I, R and IR. We also mentioned the number of trainable parameters per CNN in Fig. 1(b), which ranges from about 11 million to 46 million. Analyzing Figs. 1(b) and 3(b-d) simultaneously, we see that there is neither a positive nor a negative linear correlation between the number of trainable parameters and the accuracy of the prediction of the IQ. Instead, we see in Fig. 3(b-d) that 2D-ResNet18 performed the overall best among 2D CNNs and 3D-ResNet18 performed the overall best among 3D CNNs in PIQ, VIQ, and FSIQ prediction tasks.

As we mentioned in section that several studies used sMRI-based regional brain volumes as features in different machine learning methods to predict intelligence scores^4^,6–22,27–31. These studies used ∼8,500 healthy subjects for model training and then predicted the residual PIQ score of more than 3,500 adolescents with a mean square error (MSE) ranging from 86 to 103 (for a range of true residual PIQ score of [−40, 30]), or a correlation of 10% (*p* < 0.05). On the other hand, despite our dataset being much smaller (i.e., 850 subjects) than that in state of the art, our study reported an average mean absolute error in the range of 11.0–11.5 (see supplementary Fig. 1 and 2), which is not very different statistically than the previous studies.

Furthermore, several studies^2^,^3,32^ predicted the FSIQ score, which showed a correlation of 30-70% (*p* < 0.01) between the ground truth and estimated absolute FSIQ scores. These studies used a dataset of size less than 250 healthy subjects with an age distribution of 6–27 years. On the other hand, our dataset consists of 850 healthy (51%) and autistic (49%) subjects with an age distribution of 6–64 years. Therefore, our study-found correlation (*r*) of ∼15% between the ground truth and estimated absolute FSIQ scores (see Figs. 2(a, c) and 3(a)) cannot be fairly compared to that of the state of the art.

Our study demonstrated the potential of predicting absolute and residual PIQ, VIQ, and FSIQ scores directly from the unsegmented structural brain MRIs of 6-64 years old healthy and autistic subjects using 2D and 3D deep CNNs. However, the prediction performance in terms of Pearson correlation coefficient (*r*) and mean absolute error is weak. This scenario yields several open research questions:

- *Are 850 training samples sufficient?* We mentioned in Fig. 1(b) that the number of trainable parameters in the CNNs used in this study is more than 10 million. These models usually require a large number of datasets to get optimally trained. For example, 2D-ResNet18 and 2D-VGG8 showed human-like classification performance when trained on a 1 million ImageNet computer vision dataset. On the other hand, we cannot produce augmented data from our base 850 subjects as augmenting original structural MRI images would distort the spatial information, disentangling the relation between IQ and brain anatomy. Therefore, it is necessary to use a much larger dataset (i.e., ≫ 850) in the models used in this study to check the effect of the sample size on deep CNN-based IQ prediction.
- *Are state-of-the-art CNNs able to capture the IQ-specific discriminatory features in the structural brain MRI?* In a classical CNN like the type we used in this study, the learned features in the first layer capture low-level features (e.g., edges), the second layer detects motifs by spotting particular arrangements of edges, the third layer assembles motifs into larger combinations representing parts of objects, and subsequent layers detect objects as combinations of this parts^33^. These features of a classical CNN tend to ignore diffuse textural features^34^–36 that are often important for medical imaging applications. This inability to learn diffused textural features from medical images by conventional CNNs may also negatively affect IQ prediction tasks. Thus, further study in this avenue is also necessary to fully utilize the power of deep CNNs in IQ prediction tasks.
- *Are demographic and diagnostic covariates necessary to be included in an IQ prediction model?* We observed in Figs. 2 and 3(a) that absolute IQ prediction performance in terms of mean correlation coefficient and the mean absolute error are better than that for residual IQ scores. Residual IQ scores are estimated by removing the effect of demographic information such as age, sex, race, and socio-economic status and diagnostic information from the absolute IQ scores (see Eq. 1 in our case). Since removing the effect of demographic and diagnosis information from the absolute IQ scores results in deteriorated prediction performance, further investigation is necessary to elucidate the effect of demographic and diagnostic background on IQ scores.
- *How significant is the relation between brain size and IQ?* In this study, we used three types of input to CNNs, namely T1-weighted intensity (i.e., I), RAVENS map (i.e., R), and a combination of I and R (i.e., IR). An intensity map contains spatially localized contrast and textural intensity information of the brain. On the other hand, the RAVENS map contains morphometry information in terms of voxel value that indicates the degree of volumetric deformation required for a particular voxel in a brain to be registered to the brain atlas. Thus, the RAVENS map provides information on the relative size of a brain compared to the atlas, while intensity-based information is completely disentangled from information related to brain size. We see in this study that the absolute and residual IQ prediction performance is overall better for the contrast map than for the RAVENS map (for example, see Figs. 2(c, d) and 3). Thus, it is also necessary to investigate the effect of brain size in deep CNN-based absolute and residual IQ prediction tasks.

## Methods

### Data

We accessed 1,085 T1-weighted MRI scans from Autism Brain Imaging Data Exchange (ABIDE I)^37^ and included 850 scans in this study due to the availability of the non-zero PIQ, VIQ, and FSIQ scores (see Fig. 1(a)). Subjects’ ages range from 6-64 years (mean 16.79±7.28 years). The numbers of men and women were 725 (85%) and 125 (15%), respectively. Furthermore, 417 (49%) were autistic patients and the rest (51%) were healthy controls among these subjects. These data are collected in 15 North American and European sites. The ground truth FSIQ, PIQ, and VIQ scores also came with the sMRI data.

We used a pipeline of pre-processing operations to harmonize the data. The preprocessing steps involved (i) N4 bias correction^38^, (ii) field-of-view normalization^39^, (iii) multi-atlas skull stripping^40^, (iv) skull-stripped T1-weighted MRI images are affine registered to the SRI atlas^41^ by deformable registration via attribute matching and mutual-saliency (DRAMMS) algorithms^42^, and (v) splitting the registered brain volumes into intensity images containing contrast information and RAVENS map^24^. Then we cropped 3D brain volumes to a size of 130×170×140 voxels by removing background empty spaces around the brain. We also manually checked all MRI scans to remove failed MRIs with severe artifacts or poor registration.

### Residual Intelligence Score

To estimate residual FSIQ, PIQ, and VIQ scores, we used age, sex, diagnostic group, and data collection site as independent variables and FSIQ or PIQ or VIQ as the dependent variable:

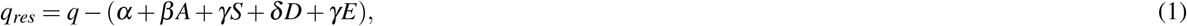

where *q*_*res*_ and *q* are residual and ground truth scores of the FSIQ/PIQ/VIQ scores, respectively, *A* denotes age in years, *S* denotes sex (1: male, 2: female), *D* denotes the diagnostic group (1: healthy, 2: autistic), *E* : (1 ≤ *E* ≤ 15) denotes the sample collection sites, and *α, β, γ, δ*, and *γ* are parameters of linear regression.

### Deep Convolutional Neural Networks

To predict actual and residual FSIQ, PIQ, and VIQ scores from brain sMRI, we implemented two 2D deep CNN models, namely 2D-ResNet18^43^ and 2D-VGG8^44^, and four 3D deep CNN models, namely 3D-ResNet18 and 3D-ResNet50^43^, and 3D-DenseNet121 and 3D-DenseNet169^45^. The number of trainable parameters per CNN is shown in Fig. 1(b). For 2D CNNs, we used 2D axial slices of brain MRI as input with different numbers of slices (i.e., 5, 10, 20, 40, 70, 100, and 130) as image channels. On the other hand, we used the entire 3D brain volume as input to 3D CNNs. We used the mean absolute error loss function to train our CNNs defined as:

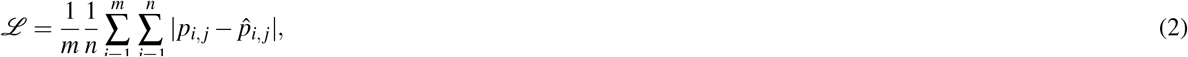

where *p* and 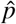 are the ground truth (actual *q* or residual *q*_*res*_) and the predicted IQ values, respectively, *m* denotes the total number of training data in a batch, and *n* denotes the number of output classes (i.e., either 1 or 3 in this study).

### Training of Deep CNNs

We chose the Adam optimizer with a learning rate of 0.01 to train all our deep CNNs. We also chose a batch size of 16 images and 4 volumes for 2D and 3D CNNs, respectively. We implemented our models in PyTorch version 1.6.0 and Python version 3.8.10. The training was performed on a workstation with an Intel E5-2650 v4 Broadwell 2.2 GHz processor, an Nvidia Titan RTX GPU with 24 GB of VRAM, and 8 GB of RAM.

