## Supplementary Tables for "Can deep learning predict human intelligence from structural brain MRI?"

### Suppementary Results

#### Absolute and Residual IQ Prediction by 2D CNNs

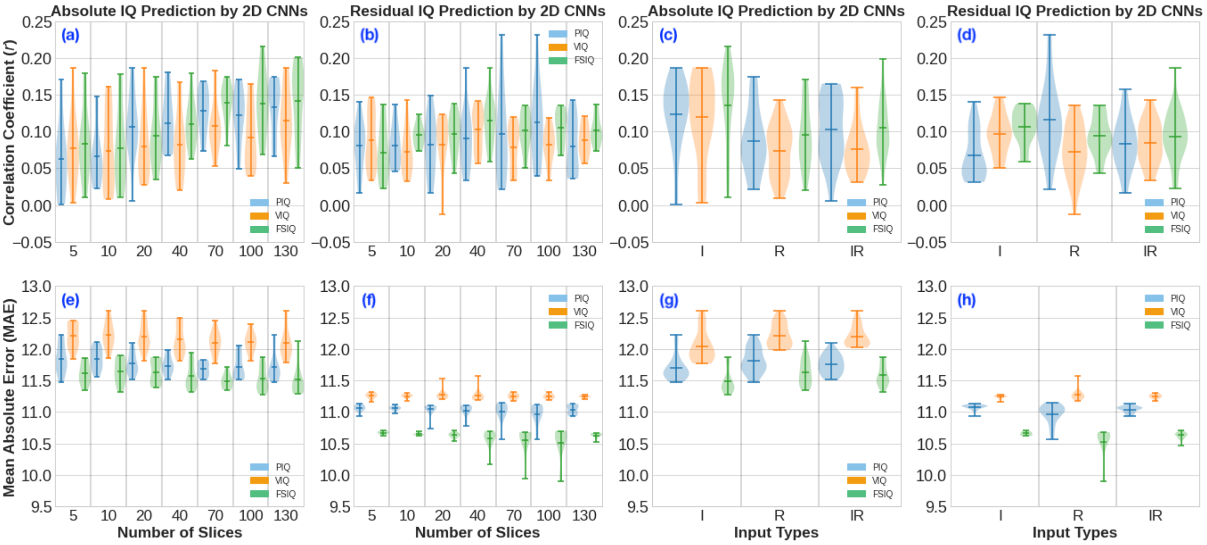

Figure 1: Violin plots showing absolute and residual IQ prediction performance in terms of Pearson correlation coefficient ($r$) (a-d) and mean absolute error (e-h) by both 2D-ResNet18 and 2D-VGG8 in settings 1 and 2. (a, e) Predicted absolute PIQ, VIQ, and FSIQ scores *vs.* numbers of slices, (b, f) predicted residual PIQ, VIQ, and FSIQ scores *vs.* numbers of slices, (c, g) predicted absolute PIQ, VIQ, and FSIQ scores *vs.* input types, and (d, h) predicted residual PIQ, VIQ, and FSIQ scores *vs.* input types.

Table 1: Absolute IQ prediction performance by 2D-ResNet18 in setting 1. MAE and correlation coefficient ($r$) are combined over 5-folds. For $p$-values equal to or smaller than 0.0001 are shown as 0.0001. Acronyms are used as, Exp: experiment number, I: intensity map as input, R: RAVENS map as input, and IR: combined intensity and RAVENS maps as input.

| **Exp.** | **IQ** | **Data** | **# of Slices** | **Mean MAE** | $r$ | $p$**-value** |
|---|---|---|---|---|---|---|
| 1 | | | 5 | 11.47±9.29 | 0.171 | 0.0001 |
| 2 | | | 10 | 11.56±9.28 | 0.148 | 0.0001 |
| 3 | | | 20 | 11.52±9.11 | 0.186 | 0.0001 |
| 4 | PIQ | I | 40 | 11.59±9.05 | 0.181 | 0.0001 |
| 5 | | | 70 | 11.51±9.22 | 0.168 | 0.0001 |
| 6 | | | 100 | 11.60±9.09 | 0.164 | 0.0001 |
| 7 | | | 130 | 11.56±9.16 | 0.162 | 0.0001 |
| 8 | | | 5 | 11.71±9.29 | 0.023 | 0.5045 |
| 9 | | | 10 | 11.75±9.27 | 0.060 | 0.0813 |
| 10 | | | 20 | 11.64±9.30 | 0.112 | 0.0010 |
| 11 | PIQ | R | 40 | 11.68±9.44 | 0.090 | 0.0085 |
| 12 | | | 70 | 11.63±9.43 | 0.112 | 0.0011 |
| 13 | | | 100 | 11.57±9.27 | 0.154 | 0.0001 |
| 14 | | | 130 | 11.47±9.51 | 0.174 | 0.0001 |
| 15 | | | 5 | 11.65±9.36 | 0.072 | 0.0353 |
| 16 | | | 10 | 11.76±9.30 | 0.058 | 0.0936 |
| 17 | | | 20 | 11.66±9.30 | 0.097 | 0.0047 |
| 18 | PIQ | IR | 40 | 11.51±9.25 | 0.162 | 0.0001 |

Continued on next page

**Table 1 – continued from previous page**

| Exp. | IQ | Data | # of Slices | Mean MAE | $r$ | $p$-value |
|---|---|---|---|---|---|---|
| 19 | | | 70 | 11.54±9.33 | 0.140 | 0.0001 |
| 20 | | | 100 | 11.51±9.43 | 0.128 | 0.0002 |
| 21 | | | 130 | 11.67±9.21 | 0.119 | 0.0005 |
| 22 | | | 5 | 11.84±9.18 | 0.186 | 0.0001 |
| 23 | | | 10 | 11.85±9.30 | 0.161 | 0.0001 |
| 24 | | | 20 | 11.82±9.19 | 0.187 | 0.0001 |
| 25 | VIQ | I | 40 | 11.89±9.26 | 0.158 | 0.0001 |
| 26 | | | 70 | 11.77±9.28 | 0.183 | 0.0001 |
| 27 | | | 100 | 11.82±9.34 | 0.157 | 0.0001 |
| 28 | | | 130 | 11.87±9.24 | 0.176 | 0.0001 |
| 29 | | | 5 | 12.06±9.25 | 0.085 | 0.0130 |
| 30 | | | 10 | 12.13±9.40 | 0.143 | 0.0001 |
| 31 | | | 20 | 12.07±9.33 | 0.060 | 0.0795 |
| 32 | VIQ | R | 40 | 11.98±9.44 | 0.090 | 0.0088 |
| 33 | | | 70 | 12.04±9.28 | 0.118 | 0.0006 |
| 34 | | | 100 | 12.04±9.48 | 0.040 | 0.2495 |
| 35 | | | 130 | 12.00±9.40 | 0.082 | 0.0169 |
| 36 | | | 5 | 12.05±9.23 | 0.105 | 0.0022 |
| 37 | | | 10 | 12.15±9.45 | 0.046 | 0.1770 |
| 38 | | | 20 | 12.11±9.27 | 0.065 | 0.0571 |
| 39 | VIQ | IR | 40 | 12.04±9.29 | 0.069 | 0.0451 |
| 40 | | | 70 | 12.06±9.23 | 0.090 | 0.0084 |
| 41 | | | 100 | 12.04±9.42 | 0.067 | 0.0497 |
| 42 | | | 130 | 12.06±9.33 | 0.069 | 0.0446 |
| 43 | | | 5 | 11.35±8.61 | 0.179 | 0.0001 |
| 44 | | | 10 | 11.32±8.62 | 0.178 | 0.0001 |
| 45 | | | 20 | 11.39±8.64 | 0.160 | 0.0001 |
| 46 | FSIQ | I | 40 | 11.37±8.61 | 0.179 | 0.0001 |
| 47 | | | 70 | 11.35±8.75 | 0.139 | 0.0001 |
| 48 | | | 100 | 11.28±8.69 | 0.176 | 0.0001 |
| 49 | | | 130 | 11.29±8.56 | 0.201 | 0.0001 |
| 50 | | | 5 | 11.49±8.79 | 0.078 | 0.0234 |
| 51 | | | 10 | 11.53±8.74 | 0.048 | 0.1624 |
| 52 | | | 20 | 11.44±8.64 | 0.136 | 0.0001 |
| 53 | FSIQ | R | 40 | 11.44±8.74 | 0.100 | 0.0035 |
| 54 | | | 70 | 11.36±8.86 | 0.163 | 0.0001 |
| 55 | | | 100 | 11.45±8.92 | 0.153 | 0.0001 |
| 56 | | | 130 | 11.38±8.77 | 0.163 | 0.0001 |
| 57 | | | 5 | 11.44±8.76 | 0.105 | 0.0022 |
| 58 | | | 10 | 11.44±8.87 | 0.122 | 0.0004 |
| 59 | | | 20 | 11.50±8.68 | 0.084 | 0.0144 |
| 60 | FSIQ | IR | 40 | 11.32±8.85 | 0.144 | 0.0001 |
| 61 | | | 70 | 11.40±8.75 | 0.145 | 0.0001 |
| 62 | | | 100 | 11.38±8.79 | 0.189 | 0.0001 |
| 63 | | | 130 | 11.38±8.85 | 0.121 | 0.0004 |

Table 2: Absolute IQ prediction performance by 2D-ResNet18 in setting 2. MAE and correlation coefficient ($r$) are combined over 5-folds. For $p$-values equal to or smaller than 0.0001 are shown as 0.0001. Acronyms are used as, Exp: experiment number, I: intensity map as input, R: RAVENS map as input, and IR: combined intensity and RAVENS maps as input.

| **Exp.** | **IQ** | **Data** | **# of Slices** | **Mean MAE** | ***r*** | ***p*-value** |
|---|---|---|---|---|---|---|
| 64 | FSIQ | I | 5 | 11.44±8.56 | 0.156 | 0.0001 |
| | PIQ | | | 11.58±9.22 | 0.131 | 0.0001 |
| | VIQ | | | 12.00±9.20 | 0.126 | 0.0002 |
| 65 | FSIQ | I | 10 | 11.42±8.54 | 0.166 | 0.0001 |
| | PIQ | | | 11.65±9.12 | 0.137 | 0.0001 |
| | VIQ | | | 12.00±9.34 | 0.100 | 0.0034 |
| 66 | FSIQ | I | 20 | 11.41±8.56 | 0.174 | 0.0001 |
| | PIQ | | | 11.61±9.15 | 0.154 | 0.0001 |
| | VIQ | | | 12.04±9.18 | 0.133 | 0.0001 |
| 67 | FSIQ | I | 40 | 11.46±8.59 | 0.135 | 0.0001 |
| | PIQ | | | 11.67±9.18 | 0.110 | 0.0014 |
| | VIQ | | | 12.00±9.24 | 0.108 | 0.0015 |
| 68 | FSIQ | I | 70 | 11.35±8.60 | 0.174 | 0.0001 |
| | PIQ | | | 11.58±9.28 | 0.132 | 0.0001 |
| | VIQ | | | 12.00±9.28 | 0.112 | 0.0010 |
| 69 | FSIQ | I | 100 | 11.38±8.63 | 0.155 | 0.0001 |
| | PIQ | | | 11.64±9.17 | 0.127 | 0.0002 |
| | VIQ | | | 11.99±9.23 | 0.121 | 0.0004 |
| 70 | FSIQ | I | 130 | 11.35±8.59 | 0.178 | 0.0001 |
| | PIQ | | | 11.58±9.19 | 0.142 | 0.0001 |
| | VIQ | | | 11.90±9.29 | 0.136 | 0.0001 |
| 71 | FSIQ | R | 5 | 11.59±8.79 | 0.097 | 0.0045 |
| | PIQ | | | 11.85±9.51 | 0.073 | 0.0333 |
| | VIQ | | | 12.12±9.39 | 0.069 | 0.0431 |
| 72 | FSIQ | R | 10 | 11.63±8.82 | 0.020 | 0.5540 |
| | PIQ | | | 11.78±9.33 | 0.022 | 0.5155 |
| | VIQ | | | 12.32±9.37 | 0.009 | 0.8025 |
| 73 | FSIQ | R | 20 | 11.53±8.75 | 0.089 | 0.0097 |
| | PIQ | | | 11.71±9.26 | 0.091 | 0.0078 |
| | VIQ | | | 12.08±9.52 | 0.032 | 0.3459 |
| 74 | FSIQ | R | 40 | 11.46±8.72 | 0.096 | 0.0053 |
| | PIQ | | | 11.74±9.38 | 0.068 | 0.0482 |
| | VIQ | | | 11.99±9.38 | 0.086 | 0.0120 |
| 75 | FSIQ | R | 70 | 11.46±8.70 | 0.106 | 0.0020 |
| | PIQ | | | 11.65±9.35 | 0.073 | 0.0322 |
| | VIQ | | | 12.00±9.28 | 0.095 | 0.0054 |
| 76 | FSIQ | R | 100 | 11.36±8.77 | 0.130 | 0.0001 |
| | PIQ | | | 11.53±9.35 | 0.131 | 0.0001 |
| | VIQ | | | 12.13±9.40 | 0.045 | 0.1895 |
| 77 | FSIQ | R | 130 | 11.35±8.70 | 0.171 | 0.0001 |
| | PIQ | | | 11.59±9.25 | 0.167 | 0.0001 |
| | VIQ | | | 11.99±9.40 | 0.118 | 0.0006 |
| 78 | FSIQ | IR | 5 | 11.54±8.65 | 0.082 | 0.0166 |
| | PIQ | | | 11.78±9.24 | 0.046 | 0.1827 |
| | VIQ | | | 12.13±9.16 | 0.076 | 0.0258 |
| 79 | FSIQ | IR | 10 | 11.56±8.70 | 0.070 | 0.0412 |
| | PIQ | | | 11.67±9.30 | 0.068 | 0.0486 |
| | VIQ | | | 12.21±9.29 | 0.039 | 0.2523 |

Continued on next page

**Table 2 – continued from previous page**

| Exp. | IQ | Data | # of Slices | Mean MAE | $r$ | $p$-value |
|---|---|---|---|---|---|---|
| 80 | FSIQ<br>PIQ<br>VIQ | IR | 20 | 11.51±8.69<br>11.69±9.29<br>12.05±9.32 | 0.102<br>0.104<br>0.086 | 0.0030<br>0.0024<br>0.0121 |
| 81 | FSIQ<br>PIQ<br>VIQ | IR | 40 | 11.45±8.65<br>11.60±9.38<br>12.08±9.25 | 0.123<br>0.119<br>0.086 | 0.0003<br>0.0005<br>0.0125 |
| 82 | FSIQ<br>PIQ<br>VIQ | IR | 70 | 11.50±8.84<br>11.58±9.43<br>12.16±9.31 | 0.081<br>0.084<br>0.053 | 0.0180<br>0.0140<br>0.1218 |
| 83 | FSIQ<br>PIQ<br>VIQ | IR | 100 | 11.47±8.60<br>11.55±9.23<br>12.11±9.31 | 0.140<br>0.146<br>0.063 | 0.0001<br>0.0001<br>0.0666 |
| 84 | FSIQ<br>PIQ<br>VIQ | IR | 130 | 11.50±8.69<br>11.63±9.33<br>12.08±9.20 | 0.091<br>0.087<br>0.092 | 0.0078<br>0.0115<br>0.0070 |

Table 3: Absolute IQ prediction performance by 2D-VGG8 in setting 1. MAE and correlation coefficient ($r$) are combined over 5-folds. For $p$-values equal to or smaller than 0.0001 are shown as 0.0001. Acronyms are used as, Exp: experiment number, I: intensity map as input, R: RAVENS map as input, and IR: combined intensity and RAVENS maps as input.

| Exp. | IQ | Data | # of Slices | Mean MAE | $r$ | $p$-value |
|---|---|---|---|---|---|---|
| 85 | PIQ | I | 5 | 12.23±9.78 | 0.005 | 0.8890 |
| 86 | | | 10 | 11.90±9.43 | 0.050 | 0.1452 |
| 87 | | | 20 | 11.75±9.24 | 0.118 | 0.0006 |
| 88 | | | 40 | 11.72±9.17 | 0.116 | 0.0007 |
| 89 | | | 70 | 11.73±9.26 | 0.113 | 0.0010 |
| 90 | | | 100 | 11.55±9.24 | 0.171 | 0.0001 |
| 91 | | | 130 | 11.61±9.18 | 0.150 | 0.0001 |
| 92 | PIQ | R | 5 | 11.80±9.41 | 0.077 | 0.0252 |
| 93 | | | 10 | 12.01±9.51 | 0.038 | 0.2717 |
| 94 | | | 20 | 11.93±9.38 | 0.083 | 0.0152 |
| 95 | | | 40 | 11.75±9.28 | 0.121 | 0.0004 |
| 96 | | | 70 | 11.81±9.47 | 0.111 | 0.0011 |
| 97 | | | 100 | 12.06±9.37 | 0.073 | 0.0325 |
| 98 | | | 130 | 12.22±9.58 | 0.066 | 0.0532 |
| 99 | PIQ | IR | 5 | 11.96±9.65 | 0.063 | 0.0671 |
| 100 | | | 10 | 11.95±9.47 | 0.035 | 0.3136 |
| 101 | | | 20 | 11.74±9.31 | 0.158 | 0.0001 |
| 102 | | | 40 | 11.93±9.33 | 0.149 | 0.0001 |
| 103 | | | 70 | 11.81±9.30 | 0.162 | 0.0001 |
| 104 | | | 100 | 11.87±9.41 | 0.123 | 0.0003 |
| 105 | | | 130 | 11.81±9.24 | 0.155 | 0.0001 |
| 106 | VIQ | I | 5 | 12.44±9.69 | 0.047 | 0.1752 |
| 107 | | | 10 | 12.45±9.29 | 0.023 | 0.5007 |
| 108 | | | 20 | 12.16±9.61 | 0.098 | 0.0044 |
| 109 | | | 40 | 11.82±9.33 | 0.167 | 0.0001 |
| 110 | | | 70 | 11.84±9.44 | 0.163 | 0.0001 |
| 111 | | | 100 | 11.90±9.42 | 0.165 | 0.0001 |

Continued on next page

**Table 3 – continued from previous page**

| Exp. | IQ | Data | # of Slices | Mean MAE | *r* | *p*-value |
|---|---|---|---|---|---|---|
| 112 | | | 130 | 11.79±9.27 | 0.186 | 0.0001 |
| 113 | | | 5 | 12.23±9.67 | 0.094 | 0.0061 |
| 114 | | | 10 | 12.21±9.45 | 0.094 | 0.0059 |
| 115 | | | 20 | 12.17±9.27 | 0.123 | 0.0003 |
| 116 | VIQ | R | 40 | 12.50±9.49 | 0.020 | 0.5576 |
| 117 | | | 70 | 12.22±9.73 | 0.099 | 0.0040 |
| 118 | | | 100 | 12.27±9.61 | 0.104 | 0.0025 |
| 119 | | | 130 | 12.49±9.53 | 0.113 | 0.0009 |
| 120 | | | 5 | 12.33±9.46 | 0.040 | 0.2497 |
| 121 | | | 10 | 12.02±9.32 | 0.147 | 0.0001 |
| 122 | | | 20 | 12.37±9.43 | 0.043 | 0.2135 |
| 123 | VIQ | IR | 40 | 12.44±9.82 | 0.031 | 0.3741 |
| 124 | | | 70 | 12.29±9.49 | 0.073 | 0.0345 |
| 125 | | | 100 | 12.19±9.31 | 0.129 | 0.0002 |
| 126 | | | 130 | 12.23±9.48 | 0.100 | 0.0035 |
| 127 | | | 5 | 11.70±8.96 | 0.060 | 0.0782 |
| 128 | | | 10 | 11.85±8.98 | 0.011 | 0.7547 |
| 129 | | | 20 | 11.78±8.93 | 0.035 | 0.3090 |
| 130 | FSIQ | I | 40 | 11.51±8.71 | 0.106 | 0.0019 |
| 131 | | | 70 | 11.51±8.53 | 0.160 | 0.0001 |
| 132 | | | 100 | 11.37±8.79 | 0.216 | 0.0001 |
| 133 | | | 130 | 11.40±8.74 | 0.165 | 0.0001 |
| 134 | | | 5 | 11.62±9.00 | 0.074 | 0.0307 |
| 135 | | | 10 | 11.90±8.94 | 0.036 | 0.2924 |
| 136 | | | 20 | 11.68±8.94 | 0.074 | 0.0309 |
| 137 | FSIQ | R | 40 | 11.94±9.12 | 0.075 | 0.0294 |
| 138 | | | 70 | 11.62±8.79 | 0.122 | 0.0004 |
| 139 | | | 100 | 11.67±8.92 | 0.120 | 0.0005 |
| 140 | | | 130 | 11.90±9.08 | 0.080 | 0.0203 |
| 141 | | | 5 | 11.81±9.11 | 0.028 | 0.4096 |
| 142 | | | 10 | 11.70±8.83 | 0.089 | 0.0093 |
| 143 | | | 20 | 11.85±9.09 | 0.062 | 0.0697 |
| 144 | FSIQ | IR | 40 | 11.80±8.94 | 0.096 | 0.0052 |
| 145 | | | 70 | 11.51±9.00 | 0.128 | 0.0002 |
| 146 | | | 100 | 11.76±8.87 | 0.093 | 0.0064 |
| 147 | | | 130 | 11.58±8.80 | 0.124 | 0.0003 |

Table 4: Absolute IQ prediction performance by 2D-VGG8 in setting 2. MAE and correlation coefficient ($r$) are combined over 5-folds. For $p$-values equal to or smaller than 0.0001 are shown as 0.0001. Acronyms are used as, Exp: experiment number, I: intensity map as input, R: RAVENS map as input, and IR: combined intensity and RAVENS maps as input.

| Exp. | IQ | Data | # of Slices | Mean MAE | *r* | *p*-value |
|---|---|---|---|---|---|---|
| | FSIQ | | | 11.84±8.89 | 0.010 | 0.7643 |
| 148 | PIQ | I | 5 | 12.14±9.58 | 0.001 | 0.9795 |
| | VIQ | | | 12.45±9.74 | 0.003 | 0.9397 |
| | FSIQ | | | 11.87±8.91 | 0.069 | 0.0432 |
| 149 | PIQ | I | 10 | 12.11±9.35 | 0.086 | 0.0123 |
| | VIQ | | | 12.60±9.73 | 0.008 | 0.8251 |
| | FSIQ | | | 11.84±8.74 | 0.072 | 0.0357 |

Continued on next page

**Table 4 – continued from previous page**

| Exp. | IQ | Data | # of Slices | Mean MAE | $r$ | $p$-value |
|---|---|---|---|---|---|---|
| 150 | PIQ | I | 20 | 11.91±9.35 | 0.078 | 0.0235 |
| | VIQ | | | 12.43±9.54 | 0.027 | 0.4289 |
| | FSIQ | | | 11.49±8.70 | 0.119 | 0.0005 |
| 151 | PIQ | I | 40 | 11.78±9.33 | 0.079 | 0.0213 |
| | VIQ | | | 12.16±9.32 | 0.084 | 0.0142 |
| | FSIQ | | | 11.47±8.78 | 0.146 | 0.0001 |
| 152 | PIQ | I | 70 | 11.70±9.16 | 0.147 | 0.0001 |
| | VIQ | | | 12.17±9.31 | 0.108 | 0.0017 |
| | FSIQ | | | 11.54±8.86 | 0.125 | 0.0003 |
| 153 | PIQ | I | 100 | 11.74±9.43 | 0.111 | 0.0012 |
| | VIQ | | | 12.14±9.57 | 0.097 | 0.0046 |
| | FSIQ | | | 11.36±8.75 | 0.161 | 0.0001 |
| 154 | PIQ | I | 130 | 11.62±9.31 | 0.132 | 0.0001 |
| | VIQ | | | 12.10±9.24 | 0.123 | 0.0003 |
| | FSIQ | | | 11.86±8.89 | 0.050 | 0.1447 |
| 155 | PIQ | R | 5 | 12.17±9.33 | 0.021 | 0.5488 |
| | VIQ | | | 12.41±9.48 | 0.038 | 0.2661 |
| | FSIQ | | | 11.81±9.10 | 0.045 | 0.1874 |
| 156 | PIQ | R | 10 | 12.07±9.51 | 0.032 | 0.3504 |
| | VIQ | | | 12.44±9.50 | 0.035 | 0.3055 |
| | FSIQ | | | 11.83±8.95 | 0.093 | 0.0068 |
| 157 | PIQ | R | 20 | 11.97±9.33 | 0.094 | 0.0063 |
| | VIQ | | | 12.42±9.46 | 0.058 | 0.0927 |
| | FSIQ | | | 11.83±9.05 | 0.063 | 0.0657 |
| 158 | PIQ | R | 40 | 11.98±9.54 | 0.068 | 0.0479 |
| | VIQ | | | 12.38±9.64 | 0.048 | 0.1591 |
| | FSIQ | | | 11.71±8.81 | 0.155 | 0.0001 |
| 159 | PIQ | R | 70 | 11.78±9.37 | 0.163 | 0.0001 |
| | VIQ | | | 12.45±9.49 | 0.091 | 0.0077 |
| | FSIQ | | | 11.76±8.98 | 0.069 | 0.0446 |
| 160 | PIQ | R | 100 | 12.00±9.55 | 0.049 | 0.1568 |
| | VIQ | | | 12.23±9.63 | 0.055 | 0.1061 |
| | FSIQ | | | 12.12±9.30 | 0.051 | 0.1367 |
| 161 | PIQ | R | 130 | 12.10±9.49 | 0.075 | 0.0297 |
| | VIQ | | | 12.60±9.85 | 0.030 | 0.3900 |
| | FSIQ | | | 11.64±9.09 | 0.080 | 0.0203 |
| 162 | PIQ | IR | 5 | 11.81±9.58 | 0.071 | 0.0396 |
| | VIQ | | | 12.44±9.50 | 0.061 | 0.0750 |
| | FSIQ | | | 11.64±8.92 | 0.080 | 0.0201 |
| 163 | PIQ | IR | 10 | 11.95±9.37 | 0.066 | 0.0545 |
| | VIQ | | | 12.23±9.43 | 0.075 | 0.0291 |
| | FSIQ | | | 11.87±8.95 | 0.053 | 0.1236 |
| 164 | PIQ | IR | 20 | 12.09±9.57 | 0.006 | 0.8540 |
| | VIQ | | | 12.60±9.29 | 0.047 | 0.1743 |
| | FSIQ | | | 11.72±8.75 | 0.088 | 0.0103 |
| 165 | PIQ | IR | 40 | 11.87±9.30 | 0.073 | 0.0337 |
| | VIQ | | | 12.49±9.25 | 0.039 | 0.2541 |
| | FSIQ | | | 11.59±8.82 | 0.152 | 0.0001 |
| 166 | PIQ | IR | 70 | 11.83±9.32 | 0.138 | 0.0001 |
| | VIQ | | | 12.20±9.50 | 0.112 | 0.0011 |
| | FSIQ | | | 11.87±8.94 | 0.083 | 0.0159 |
| 167 | PIQ | IR | 100 | 11.94±9.41 | 0.089 | 0.0098 |
| | VIQ | | | 12.40±9.74 | 0.058 | 0.0928 |
| | FSIQ | | | 11.51±8.67 | 0.199 | 0.0001 |

Continued on next page

**Table 4 – continued from previous page**

| Exp. | IQ | Data | # of Slices | Mean MAE | $r$ | $p$-value |
|---|---|---|---|---|---|---|
| 168 | PIQ | IR | 130 | 11.73±9.42 | 0.165 | 0.0001 |
| | VIQ | | | 12.03±9.50 | 0.160 | 0.0001 |

Table 5: Residual IQ prediction performance by 2D-ResNet18 in setting 1. MAE and correlation coefficient ($r$) are combined over 5-folds. For $p$-values equal to or smaller than 0.0001 are shown as 0.0001. Acronyms are used as, Exp: experiment number, I: intensity map as input, R: RAVENS map as input, and IR: combined intensity and RAVENS maps as input.

| Exp. | IQ | Data | # of Slices | Mean MAE | $r$ | $p$-value |
|---|---|---|---|---|---|---|
| 169 | | | 5 | 10.94±9.05 | 0.140 | 0.0001 |
| 170 | | | 10 | 11.02±8.99 | 0.119 | 0.0005 |
| 171 | | | 20 | 11.06±8.94 | 0.115 | 0.0008 |
| 172 | PIQ | I | 40 | 11.05±8.96 | 0.114 | 0.0008 |
| 173 | | | 70 | 11.07±9.01 | 0.076 | 0.0258 |
| 174 | | | 100 | 11.07±9.01 | 0.088 | 0.0101 |
| 175 | | | 130 | 11.09±9.03 | 0.036 | 0.2996 |
| 176 | | | 5 | 11.05±8.98 | 0.105 | 0.0021 |
| 177 | | | 10 | 10.98±8.97 | 0.137 | 0.0001 |
| 178 | | | 20 | 11.08±9.01 | 0.065 | 0.0565 |
| 179 | PIQ | R | 40 | 11.08±8.96 | 0.110 | 0.0014 |
| 180 | | | 70 | 10.94±9.02 | 0.138 | 0.0001 |
| 181 | | | 100 | 10.91±8.99 | 0.162 | 0.0001 |
| 182 | | | 130 | 10.96±9.07 | 0.117 | 0.0007 |
| 183 | | | 5 | 11.06±9.01 | 0.088 | 0.0104 |
| 184 | | | 10 | 11.04±9.23 | 0.083 | 0.0159 |
| 185 | | | 20 | 11.08±9.13 | 0.043 | 0.2068 |
| 186 | PIQ | IR | 40 | 11.04±9.14 | 0.063 | 0.0674 |
| 187 | | | 70 | 11.02±9.04 | 0.106 | 0.0019 |
| 188 | | | 100 | 10.94±9.07 | 0.157 | 0.0001 |
| 189 | | | 130 | 10.93±9.09 | 0.143 | 0.0001 |
| 190 | | | 5 | 11.16±8.99 | 0.146 | 0.0001 |
| 191 | | | 10 | 11.23±9.03 | 0.107 | 0.0018 |
| 192 | | | 20 | 11.25±9.03 | 0.098 | 0.0042 |
| 193 | VIQ | I | 40 | 11.19±9.00 | 0.141 | 0.0001 |
| 194 | | | 70 | 11.24±9.02 | 0.095 | 0.0057 |
| 195 | | | 100 | 11.19±9.04 | 0.119 | 0.0005 |
| 196 | | | 130 | 11.24±8.97 | 0.121 | 0.0004 |
| 197 | | | 5 | 11.18±8.99 | 0.136 | 0.0001 |
| 198 | | | 10 | 11.22±9.08 | 0.085 | 0.0135 |
| 199 | | | 20 | 11.21±9.06 | 0.107 | 0.0018 |
| 200 | VIQ | R | 40 | 11.21±9.03 | 0.114 | 0.0009 |
| 201 | | | 70 | 11.21±9.00 | 0.120 | 0.0004 |
| 202 | | | 100 | 11.24±9.10 | 0.082 | 0.0167 |
| 203 | | | 130 | 11.25±9.00 | 0.105 | 0.0021 |
| 204 | | | 5 | 11.24±8.98 | 0.108 | 0.0016 |
| 205 | | | 10 | 11.17±8.97 | 0.143 | 0.0001 |
| 206 | | | 20 | 11.26±8.97 | 0.123 | 0.0003 |
| 207 | VIQ | IR | 40 | 11.24±9.00 | 0.120 | 0.0004 |
| 208 | | | 70 | 11.25±9.09 | 0.052 | 0.1320 |
| 209 | | | 100 | 11.22±9.12 | 0.094 | 0.0062 |

Continued on next page

Table 5 – continued from previous page

| Exp. | IQ | Data | # of Slices | Mean MAE | $r$ | $p$-value |
|---|---|---|---|---|---|---|
| 210 | | | 130 | 11.21±9.09 | 0.078 | 0.0231 |
| 211 | | | 5 | 10.63±8.41 | 0.128 | 0.0002 |
| 212 | | | 10 | 10.64±8.40 | 0.123 | 0.0003 |
| 213 | | | 20 | 10.62±8.35 | 0.138 | 0.0001 |
| 214 | FSIQ | I | 40 | 10.64±8.39 | 0.127 | 0.0002 |
| 215 | | | 70 | 10.63±8.43 | 0.129 | 0.0002 |
| 216 | | | 100 | 10.62±8.47 | 0.099 | 0.0037 |
| 217 | | | 130 | 10.63±8.40 | 0.137 | 0.0001 |
| 218 | | | 5 | 10.68±8.50 | 0.059 | 0.0877 |
| 219 | | | 10 | 10.64±8.48 | 0.081 | 0.0182 |
| 220 | | | 20 | 10.59±8.54 | 0.098 | 0.0044 |
| 221 | FSIQ | R | 40 | 10.62±8.49 | 0.099 | 0.0040 |
| 222 | | | 70 | 10.63±8.50 | 0.084 | 0.0149 |
| 223 | | | 100 | 10.62±8.51 | 0.078 | 0.0227 |
| 224 | | | 130 | 10.63±8.48 | 0.104 | 0.0024 |
| 225 | | | 5 | 10.63±8.51 | 0.085 | 0.0131 |
| 226 | | | 10 | 10.64±8.48 | 0.089 | 0.0093 |
| 227 | | | 20 | 10.67±8.45 | 0.080 | 0.0199 |
| 228 | FSIQ | IR | 40 | 10.47±8.39 | 0.186 | 0.0001 |
| 229 | | | 70 | 10.67±8.46 | 0.066 | 0.0539 |
| 230 | | | 100 | 10.60±8.57 | 0.081 | 0.0183 |
| 231 | | | 130 | 10.65±8.49 | 0.086 | 0.0124 |

Table 6: Residual IQ prediction performance by 2D-ResNet18 in setting 2. MAE and correlation coefficient $r$ are averaged over 5-folds. For $p$-values equal to or smaller than 0.0001 are shown as 0.0001. Acronyms are used as, Exp: experiment number, I: intensity map as input, R: RAVENS map as input, and IR: combined intensity and RAVENS maps as input.

| Exp. | IQ | Data | # of Slices | Mean MAE | $r$ | $p$-value |
|---|---|---|---|---|---|---|
| | FSIQ | | | 10.63±8.37 | 0.137 | 0.0001 |
| 232 | PIQ | I | 5 | 11.12±8.90 | 0.113 | 0.0009 |
| | VIQ | | | 11.24±8.92 | 0.132 | 0.0001 |
| | FSIQ | | | 10.68±8.40 | 0.100 | 0.0036 |
| 233 | PIQ | I | 10 | 11.07±9.04 | 0.067 | 0.0525 |
| | VIQ | | | 11.27±9.05 | 0.055 | 0.1086 |
| | FSIQ | | | 10.64±8.38 | 0.122 | 0.0004 |
| 234 | PIQ | I | 20 | 11.06±8.98 | 0.100 | 0.0037 |
| | VIQ | | | 11.25±8.96 | 0.113 | 0.0010 |
| | FSIQ | | | 10.70±8.39 | 0.091 | 0.0078 |
| 235 | PIQ | I | 40 | 11.09±9.02 | 0.046 | 0.1783 |
| | VIQ | | | 11.28±9.03 | 0.063 | 0.0679 |
| | FSIQ | | | 10.67±8.41 | 0.099 | 0.0038 |
| 236 | PIQ | I | 70 | 11.11±9.00 | 0.041 | 0.2365 |
| | VIQ | | | 11.26±9.00 | 0.097 | 0.0045 |
| | FSIQ | | | 10.70±8.44 | 0.067 | 0.0494 |
| 237 | PIQ | I | 100 | 11.11±9.03 | 0.039 | 0.2575 |
| | VIQ | | | 11.21±9.09 | 0.081 | 0.0185 |
| | FSIQ | | | 10.67±8.43 | 0.088 | 0.0102 |
| 238 | PIQ | I | 130 | 11.13±8.99 | 0.039 | 0.2599 |

Continued on next page

**Table 6 – continued from previous page**

| Exp. | IQ | Data | # of Slices | Mean MAE | $r$ | $p$-value |
|---|---|---|---|---|---|---|
| | VIQ | | | 11.27±9.04 | 0.056 | 0.1030 |
| | FSIQ | | | 10.67±8.41 | 0.102 | 0.0030 |
| 239 | PIQ | R | 5 | 11.12±8.95 | 0.080 | 0.0191 |
| | VIQ | | | 11.28±8.98 | 0.084 | 0.0140 |
| | FSIQ | | | 10.67±8.48 | 0.086 | 0.0123 |
| 240 | PIQ | R | 10 | 11.12±9.00 | 0.051 | 0.1412 |
| | VIQ | | | 11.26±9.08 | 0.054 | 0.1147 |
| | FSIQ | | | 10.66±8.53 | 0.079 | 0.0219 |
| 241 | PIQ | R | 20 | 11.09±8.95 | 0.104 | 0.0025 |
| | VIQ | | | 11.26±9.11 | 0.015 | 0.6645 |
| | FSIQ | | | 10.63±8.37 | 0.086 | 0.0536 |
| 242 | PIQ | R | 40 | 10.78±8.97 | 0.082 | 0.0631 |
| | VIQ | | | 11.57±9.22 | 0.057 | 0.2012 |
| | FSIQ | | | 10.28±8.50 | 0.050 | 0.3554 |
| 243 | PIQ | R | 70 | 11.15±9.31 | 0.021 | 0.6998 |
| | VIQ | | | 11.29±8.79 | 0.035 | 0.5157 |
| | FSIQ | | | 09.90±8.55 | 0.119 | 0.1224 |
| 244 | PIQ | R | 100 | 10.69±9.48 | 0.127 | 0.0991 |
| | VIQ | | | 11.32±9.18 | 0.081 | 0.2932 |
| | FSIQ | | | 10.63±8.49 | 0.084 | 0.0140 |
| 245 | PIQ | R | 130 | 11.11±9.08 | 0.042 | 0.2178 |
| | VIQ | | | 11.25±9.05 | 0.074 | 0.0313 |
| | FSIQ | | | 10.71±8.48 | 0.051 | 0.1402 |
| 246 | PIQ | IR | 5 | 11.13±9.05 | 0.016 | 0.6311 |
| | VIQ | | | 11.30±9.03 | 0.056 | 0.1034 |
| | FSIQ | | | 10.63±8.45 | 0.098 | 0.0043 |
| 247 | PIQ | IR | 10 | 11.09±9.13 | 0.046 | 0.1833 |
| | VIQ | | | 11.28±9.04 | 0.063 | 0.0653 |
| | FSIQ | | | 10.59±8.46 | 0.116 | 0.0007 |
| 248 | PIQ | IR | 20 | 11.08±9.01 | 0.085 | 0.0129 |
| | VIQ | | | 11.29±8.95 | 0.109 | 0.0014 |
| | FSIQ | | | 10.56±8.56 | 0.135 | 0.0001 |
| 249 | PIQ | IR | 40 | 11.07±9.00 | 0.111 | 0.0012 |
| | VIQ | | | 11.23±8.97 | 0.129 | 0.0002 |
| | FSIQ | | | 10.67±8.40 | 0.097 | 0.0046 |
| 250 | PIQ | IR | 70 | 11.06±9.02 | 0.080 | 0.0203 |
| | VIQ | | | 11.31±9.03 | 0.033 | 0.3334 |
| | FSIQ | | | 10.65±8.43 | 0.124 | 0.0003 |
| 251 | PIQ | IR | 100 | 11.12±9.03 | 0.095 | 0.0056 |
| | VIQ | | | 11.23±9.01 | 0.107 | 0.0018 |
| | FSIQ | | | 10.65±8.40 | 0.100 | 0.0034 |
| 252 | PIQ | IR | 130 | 11.07±9.03 | 0.070 | 0.0427 |
| | VIQ | | | 11.23±9.01 | 0.103 | 0.0027 |

Table 7: Residual IQ prediction performance by 2D-VGG8 in setting 1. MAE and correlation coefficient ($r$) are combined over 5-folds. For $p$-values equal to or smaller than 0.0001 are shown as 0.0001. Acronyms are used as, Exp: experiment number, I: intensity map as input, R: RAVENS map as input, and IR: combined intensity and RAVENS maps as input.

| **Exp.** | **IQ** | **Data** | **# of Slices** | **Mean MAE** | $r$ | $p$**-value** |
|---|---|---|---|---|---|---|
| 253 | | | 5 | 11.01±9.04 | 0.097 | 0.0048 |
| 254 | | | 10 | 11.04±9.00 | 0.101 | 0.0031 |
| 255 | | | 20 | 11.09±9.03 | 0.060 | 0.0805 |
| 256 | PIQ | I | 40 | 11.09±9.02 | 0.067 | 0.0498 |
| 257 | | | 70 | 11.10±9.02 | 0.057 | 0.0942 |
| 258 | | | 100 | 11.09±9.03 | 0.046 | 0.1795 |
| 259 | | | 130 | 11.10±9.03 | 0.040 | 0.2490 |
| 260 | | | 5 | 11.04±9.04 | 0.089 | 0.0092 |
| 261 | | | 10 | 11.06±9.01 | 0.087 | 0.0108 |
| 262 | | | 20 | 11.05±8.92 | 0.125 | 0.0002 |
| 263 | PIQ | R | 40 | 11.00±9.03 | 0.111 | 0.0011 |
| 264 | | | 70 | 10.93±8.92 | 0.175 | 0.0001 |
| 265 | | | 100 | 10.98±9.00 | 0.133 | 0.0001 |
| 266 | | | 130 | 10.98±9.03 | 0.117 | 0.0006 |
| 267 | | | 5 | 11.06±9.05 | 0.064 | 0.0627 |
| 268 | | | 10 | 11.07±9.06 | 0.055 | 0.1091 |
| 269 | | | 20 | 11.09±9.08 | 0.017 | 0.6174 |
| 270 | PIQ | IR | 40 | 11.07±9.04 | 0.060 | 0.0810 |
| 271 | | | 70 | 10.95±9.07 | 0.134 | 0.0001 |
| 272 | | | 100 | 10.99±9.03 | 0.115 | 0.0008 |
| 273 | | | 130 | 10.99±9.04 | 0.120 | 0.0005 |
| 274 | | | 5 | 11.24±9.00 | 0.119 | 0.0005 |
| 275 | | | 10 | 11.25±9.04 | 0.073 | 0.0322 |
| 276 | | | 20 | 11.24±9.05 | 0.080 | 0.0203 |
| 277 | VIQ | I | 40 | 11.24±8.98 | 0.137 | 0.0001 |
| 278 | | | 70 | 11.21±9.05 | 0.101 | 0.0033 |
| 279 | | | 100 | 11.26±9.02 | 0.103 | 0.0025 |
| 280 | | | 130 | 11.26±9.04 | 0.085 | 0.0131 |
| 281 | | | 5 | 11.25±9.06 | 0.063 | 0.0644 |
| 282 | | | 10 | 11.25±9.13 | 0.032 | 0.3495 |
| 283 | | | 20 | 11.23±8.99 | 0.107 | 0.0018 |
| 284 | VIQ | R | 40 | 11.21±9.00 | 0.117 | 0.0006 |
| 285 | | | 70 | 11.22±9.05 | 0.093 | 0.0066 |
| 286 | | | 100 | 11.27±9.04 | 0.064 | 0.0628 |
| 287 | | | 130 | 11.20±9.02 | 0.120 | 0.0005 |
| 288 | | | 5 | 11.27±9.04 | 0.059 | 0.0878 |
| 289 | | | 10 | 11.25±9.05 | 0.069 | 0.0453 |
| 290 | | | 20 | 11.27±9.02 | 0.095 | 0.0054 |
| 291 | VIQ | IR | 40 | 11.22±9.08 | 0.081 | 0.0179 |
| 292 | | | 70 | 11.17±9.08 | 0.109 | 0.0015 |
| 293 | | | 100 | 11.23±9.10 | 0.071 | 0.0395 |
| 294 | | | 130 | 11.24±9.06 | 0.072 | 0.0357 |
| 295 | | | 5 | 10.67±8.42 | 0.090 | 0.0088 |
| 296 | | | 10 | 10.65±8.40 | 0.115 | 0.0008 |
| 297 | | | 20 | 10.66±8.38 | 0.122 | 0.0004 |
| 298 | FSIQ | I | 40 | 10.67±8.40 | 0.129 | 0.0002 |
| 299 | | | 70 | 10.64±8.41 | 0.108 | 0.0017 |
| 300 | | | 100 | 10.64±8.41 | 0.127 | 0.0002 |

Continued on next page

**Table 7 – continued from previous page**

| Exp. | IQ | Data | # of Slices | Mean MAE | *r* | *p*-value |
|---|---|---|---|---|---|---|
| 301 | | | 130 | 10.64±8.42 | 0.102 | 0.0028 |
| 302 | | | 5 | 10.66±8.54 | 0.043 | 0.2117 |
| 303 | | | 10 | 10.64±8.44 | 0.100 | 0.0036 |
| 304 | | | 20 | 10.64±8.54 | 0.083 | 0.0153 |
| 305 | FSIQ | R | 40 | 10.60±8.55 | 0.085 | 0.0136 |
| 306 | | | 70 | 10.55±8.55 | 0.130 | 0.0002 |
| 307 | | | 100 | 10.54±8.52 | 0.130 | 0.0001 |
| 308 | | | 130 | 10.52±8.54 | 0.123 | 0.0003 |
| 309 | | | 5 | 10.70±8.50 | 0.023 | 0.5090 |
| 310 | | | 10 | 10.64±8.47 | 0.073 | 0.0325 |
| 311 | | | 20 | 10.63±8.57 | 0.043 | 0.2136 |
| 312 | FSIQ | IR | 40 | 10.58±8.43 | 0.131 | 0.0001 |
| 313 | | | 70 | 10.59±8.45 | 0.124 | 0.0003 |
| 314 | | | 100 | 10.66±8.47 | 0.088 | 0.0102 |
| 315 | | | 130 | 10.64±8.49 | 0.092 | 0.0070 |

Table 8: Residual IQ prediction performance by 2D-VGG8 in setting 2. MAE and correlation coefficient ($r$) are combined over 5-folds. For $p$-values equal to or smaller than 0.0001 are shown as 0.0001. Acronyms are used as, Exp: experiment number, I: intensity map as input, R: RAVENS map as input, and IR: combined intensity and RAVENS maps as input.

| Exp. | IQ | Data | # of Slices | Mean MAE | *r* | *p*-value |
|---|---|---|---|---|---|---|
| | FSIQ | | | 10.69±8.44 | 0.064 | 0.0637 |
| 316 | PIQ | I | 5 | 11.11±9.01 | 0.040 | 0.2386 |
| | VIQ | | | 11.28±9.04 | 0.058 | 0.0937 |
| | FSIQ | | | 10.68±8.39 | 0.108 | 0.0015 |
| 317 | PIQ | I | 10 | 11.08±9.02 | 0.058 | 0.0891 |
| | VIQ | | | 11.27±9.00 | 0.101 | 0.0033 |
| | FSIQ | | | 10.71±8.43 | 0.061 | 0.0753 |
| 318 | PIQ | I | 20 | 11.10±9.02 | 0.032 | 0.3573 |
| | VIQ | | | 11.26±9.05 | 0.083 | 0.0160 |
| | FSIQ | | | 10.70±8.42 | 0.059 | 0.0845 |
| 319 | PIQ | I | 40 | 11.11±9.02 | 0.034 | 0.3160 |
| | VIQ | | | 11.25±9.03 | 0.087 | 0.0111 |
| | FSIQ | | | 10.68±8.39 | 0.102 | 0.0030 |
| 320 | PIQ | I | 70 | 11.11±9.02 | 0.031 | 0.3663 |
| | VIQ | | | 11.27±9.01 | 0.100 | 0.0036 |
| | FSIQ | | | 10.66±8.42 | 0.110 | 0.0014 |
| 321 | PIQ | I | 100 | 11.10±9.01 | 0.059 | 0.0845 |
| | VIQ | | | 11.26±9.06 | 0.051 | 0.1404 |
| | FSIQ | | | 10.67±8.39 | 0.101 | 0.0033 |
| 322 | PIQ | I | 130 | 11.10±9.01 | 0.051 | 0.1347 |
| | VIQ | | | 11.24±9.02 | 0.097 | 0.0044 |
| | FSIQ | | | 10.67±8.51 | 0.051 | 0.1397 |
| 323 | PIQ | R | 5 | 11.05±9.02 | 0.088 | 0.0104 |
| | VIQ | | | 11.32±8.99 | 0.061 | 0.0747 |
| | FSIQ | | | 10.64±8.49 | 0.093 | 0.0067 |
| 324 | PIQ | R | 10 | 11.07±8.97 | 0.097 | 0.0045 |
| | VIQ | | | 11.31±9.00 | 0.047 | 0.1695 |

Continued on next page

**Table 8 – continued from previous page**

| Exp. | IQ | Data | # of Slices | Mean MAE | $r$ | $p$-value |
|---|---|---|---|---|---|---|
| 325 | FSIQ | R | 20 | 10.54±8.45 | 0.106 | 0.0163 |
| | PIQ | | | 10.74±8.86 | 0.149 | 0.0007 |
| | VIQ | | | 11.53±9.37 | -0.013 | 0.7744 |
| 326 | FSIQ | R | 40 | 10.17±8.50 | 0.128 | 0.0182 |
| | PIQ | | | 10.84±9.30 | 0.187 | 0.0005 |
| | VIQ | | | 11.28±8.75 | 0.073 | 0.1794 |
| 327 | FSIQ | R | 70 | 09.94±8.65 | 0.135 | 0.0787 |
| | PIQ | | | 10.57±9.43 | 0.232 | 0.0023 |
| | VIQ | | | 11.32±9.19 | 0.033 | 0.6688 |
| 328 | FSIQ | R | 100 | 09.90±8.55 | 0.119 | 0.1224 |
| | PIQ | | | 10.69±9.48 | 0.127 | 0.0991 |
| | VIQ | | | 11.32±9.18 | 0.081 | 0.2932 |
| 329 | FSIQ | R | 130 | 10.65±8.53 | 0.073 | 0.0337 |
| | PIQ | | | 11.02±9.09 | 0.082 | 0.0169 |
| | VIQ | | | 11.27±9.03 | 0.061 | 0.0739 |
| 330 | FSIQ | IR | 5 | 10.68±8.52 | 0.022 | 0.5213 |
| | PIQ | | | 11.07±9.05 | 0.048 | 0.1633 |
| | VIQ | | | 11.30±9.04 | 0.034 | 0.3264 |
| 331 | FSIQ | IR | 10 | 10.69±8.45 | 0.076 | 0.0272 |
| | PIQ | | | 11.06±9.04 | 0.069 | 0.0429 |
| | VIQ | | | 11.28±9.05 | 0.042 | 0.2263 |
| 332 | FSIQ | IR | 20 | 10.64±8.44 | 0.108 | 0.0016 |
| | PIQ | | | 11.03±9.06 | 0.086 | 0.0126 |
| | VIQ | | | 11.26±9.04 | 0.067 | 0.0498 |
| 333 | FSIQ | IR | 40 | 10.63±8.40 | 0.122 | 0.0004 |
| | PIQ | | | 11.00±9.05 | 0.102 | 0.0030 |
| | VIQ | | | 11.20±9.02 | 0.111 | 0.0012 |
| 334 | FSIQ | IR | 70 | 10.65±8.59 | 0.089 | 0.0098 |
| | PIQ | | | 11.07±9.12 | 0.073 | 0.0340 |
| | VIQ | | | 11.27±9.05 | 0.070 | 0.0418 |
| 335 | FSIQ | IR | 100 | 10.58±8.54 | 0.109 | 0.0014 |
| | PIQ | | | 10.99±9.16 | 0.090 | 0.0088 |
| | VIQ | | | 11.28±8.97 | 0.093 | 0.0065 |
| 336 | FSIQ | IR | 130 | 10.58±8.44 | 0.127 | 0.0002 |
| | PIQ | | | 10.96±9.13 | 0.102 | 0.0029 |
| | VIQ | | | 11.26±9.02 | 0.085 | 0.0131 |

### Absolute and Residual IQ Prediction by 3D CNNs

Table 9: Absolute IQ prediction performance by 3D-ResNet18 in setting 1. MAE and correlation coefficient ($r$) are combined over 5-folds. For $p$-values equal to or smaller than 0.0001 are shown as 0.0001. Acronyms are used as, Exp: experiment number, I: intensity map as input, R: RAVENS map as input, and IR: combined intensity and RAVENS maps as input.

| Exp. | IQ | Data | Mean MAE | $r$ | $p$-value |
|---|---|---|---|---|---|

Continued on next page

**Table 9 – continued from previous page**

| Exp. | IQ | Data | Mean MAE | $r$ | $p$-value |
|---|---|---|---|---|---|
| 337 | | I | 11.96±9.60 | 0.118 | 0.0006 |
| 338 | PIQ | R | 11.79±9.95 | 0.191 | 0.0001 |
| 339 | | IR | 11.92±9.74 | 0.091 | 0.0082 |
| 340 | | I | 12.35±9.65 | 0.168 | 0.0001 |
| 341 | VIQ | R | 12.35±9.77 | 0.138 | 0.0001 |
| 342 | | IR | 12.43±9.76 | 0.111 | 0.0012 |
| 343 | | I | 11.72±9.01 | 0.141 | 0.0001 |
| 344 | FSIQ | R | 11.73±9.04 | 0.191 | 0.0001 |
| 345 | | IR | 11.67±9.12 | 0.150 | 0.0001 |

Table 10: Absolute IQ prediction performance by 3D-ResNet18 in setting 2. MAE and correlation coefficient ($r$) are combined over 5-folds. For $p$-values equal to or smaller than 0.0001 are shown as 0.0001. Acronyms are used as, Exp: experiment number, I: intensity map as input, R: RAVENS map as input, and IR: combined intensity and RAVENS maps as input.

| Exp. | IQ | Data | Mean MAE | $r$ | $p$-value |
|---|---|---|---|---|---|
| | FSIQ | | 11.76±8.97 | 0.136 | 0.0001 |
| 346 | PIQ | I | 12.05±9.69 | 0.068 | 0.0463 |
| | VIQ | | 12.38±9.63 | 0.166 | 0.0001 |
| | FSIQ | | 11.67±9.16 | 0.192 | 0.0001 |
| 347 | PIQ | R | 11.89±9.65 | 0.168 | 0.0001 |
| | VIQ | | 12.50±9.90 | 0.144 | 0.0001 |
| | FSIQ | | 11.73±9.06 | 0.118 | 0.0006 |
| 348 | PIQ | IR | 11.88±9.66 | 0.120 | 0.0005 |
| | VIQ | | 12.52±9.75 | 0.088 | 0.0099 |

Table 11: Absolute IQ prediction performance by 3D-ResNet50 in setting 1. MAE and correlation coefficient ($r$) are combined over 5-folds. For $p$-values equal to or smaller than 0.0001 are shown as 0.0001. Acronyms are used as, Exp: experiment number, I: intensity map as input, R: RAVENS map as input, and IR: combined intensity and RAVENS maps as input.

| Exp. | IQ | Data | Mean MAE | $r$ | $p$-value |
|---|---|---|---|---|---|
| 349 | | I | 11.98±9.67 | 0.092 | 0.0076 |
| 350 | PIQ | R | 11.93±9.59 | 0.122 | 0.0004 |
| 351 | | IR | 11.93±9.56 | 0.142 | 0.0001 |
| 352 | | I | 12.38±9.68 | 0.151 | 0.0001 |
| 353 | VIQ | R | 12.45±9.77 | 0.093 | 0.0065 |
| 354 | | IR | 12.42±9.90 | 0.069 | 0.0443 |
| 355 | | I | 11.66±9.11 | 0.134 | 0.0001 |
| 356 | FSIQ | R | 11.69±9.11 | 0.119 | 0.0005 |
| 357 | | IR | 11.77±9.20 | 0.064 | 0.0632 |

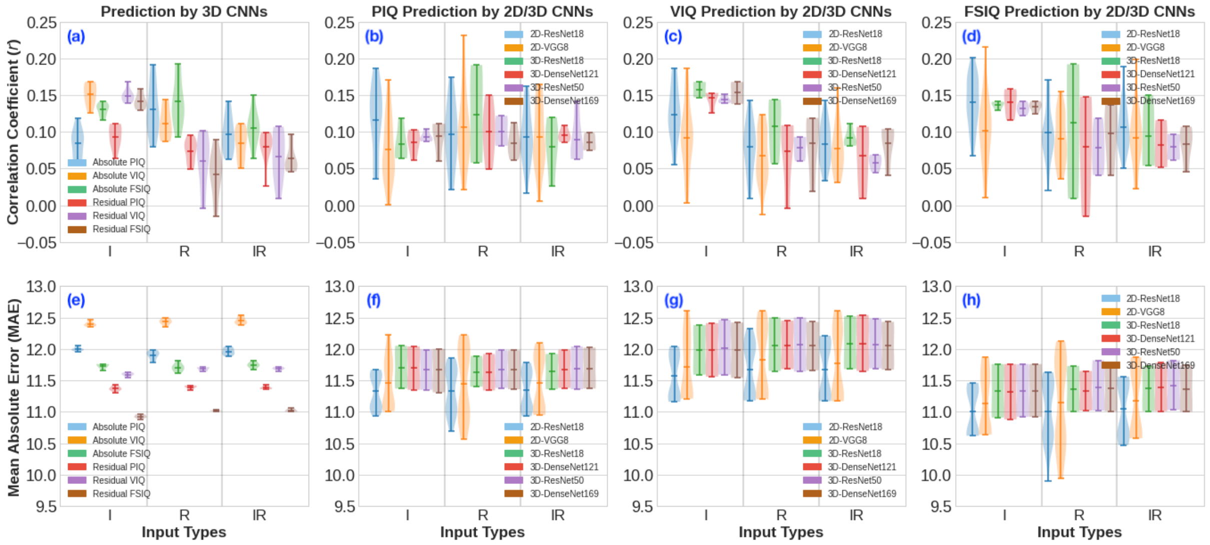


Figure 2: Violin plots showing absolute and residual IQ prediction performance in terms of Pearson correlation coefficient ($r$) and mean absolute error by 2D and 3D CNNs in settings 1 and 2. (a) Correlation and (e) mean absolute error between the ground truth and absolute/residual PIQ, VIQ, and FSIQ scores *vs.* input types by 3D-ResNet18, 3D-ResNet50, 3D-DenseNet121, and 3D-DenseNet169. (b-d) Correlation and (e-g) mean absolute error between the ground truth and actual/residual PIQ, VIQ, and FSIQ scores *vs.* input types, respectively, by different 2D and 3D CNNs.

Table 12: Absolute IQ prediction performance by 3D-ResNet50 in setting 2. MAE and correlation coefficient ($r$) are combined over 5-folds. For $p$-values equal to or smaller than 0.0001 are shown as 0.0001. Acronyms are used as, Exp: experiment number, I: intensity map as input, R: RAVENS map as input, and IR: combined intensity and RAVENS maps as input.

| Exp. | IQ | Data | Mean MAE | $r$ | $p$-value |
|---|---|---|---|---|---|
| 358 | FSIQ | I | 11.76±9.04 | 0.122 | 0.0003 |
| | PIQ | | 11.99±9.66 | 0.090 | 0.0084 |
| | VIQ | | 12.46±9.65 | 0.142 | 0.0001 |
| 359 | FSIQ | R | 11.82±9.01 | 0.093 | 0.0065 |
| | PIQ | | 11.98±9.62 | 0.115 | 0.0008 |
| | VIQ | | 12.50±9.71 | 0.093 | 0.0068 |
| 360 | FSIQ | IR | 11.82±9.07 | 0.095 | 0.0054 |
| | PIQ | | 12.04±9.60 | 0.063 | 0.0672 |
| | VIQ | | 12.48±9.85 | 0.064 | 0.0602 |

Table 13: Absolute IQ prediction performance by 3D-DenseNet121 in setting 1. MAE and correlation coefficient ($r$) are combined over 5-folds. For $p$-values equal to or smaller than 0.0001 are shown as 0.0001. Acronyms are used as, Exp: experiment number, I: intensity map as input, R: RAVENS map as input, and IR: combined intensity and RAVENS maps as input.

| Exp. | IQ | Data | Mean MAE | $r$ | $p$-value |
|---|---|---|---|---|---|

Continued on next page

Table 13 – continued from previous page

| Exp. | IQ | Data | Mean MAE | $r$ | $p$-value |
|---|---|---|---|---|---|
| 361 | | I | 11.99±9.60 | 0.092 | 0.0073 |
| 362 | PIQ | R | 11.82±9.65 | 0.150 | 0.0001 |
| 363 | | IR | 11.92±9.68 | 0.095 | 0.0055 |
| 364 | | I | 12.38±9.66 | 0.153 | 0.0001 |
| 365 | VIQ | R | 12.45±9.79 | 0.087 | 0.0116 |
| 366 | | IR | 12.41±9.80 | 0.104 | 0.0023 |
| 367 | | I | 11.72±9.08 | 0.116 | 0.0007 |
| 368 | FSIQ | R | 11.61±9.15 | 0.148 | 0.0001 |
| 369 | | IR | 11.74±9.15 | 0.103 | 0.0027 |

Table 14: Absolute IQ prediction performance by 3D-DenseNet121 in setting 2. MAE and correlation coefficient ($r$) are combined over 5-folds. For $p$-values equal to or smaller than 0.0001 are shown as 0.0001. Acronyms are used as, Exp: experiment number, I: intensity map as input, R: RAVENS map as input, and IR: combined intensity and RAVENS maps as input.

| Exp. | IQ | Data | Mean MAE | $r$ | $p$-value |
|---|---|---|---|---|---|
| | FSIQ | | 11.76±9.03 | 0.133 | 0.0001 |
| 370 | PIQ | I | 12.04±9.66 | 0.062 | 0.0704 |
| | VIQ | | 12.41±9.75 | 0.126 | 0.0002 |
| | FSIQ | | 11.65±9.16 | 0.132 | 0.0001 |
| 371 | PIQ | R | 11.92±9.67 | 0.108 | 0.0016 |
| | VIQ | | 12.40±9.86 | 0.109 | 0.0014 |
| | FSIQ | | 11.78±9.06 | 0.116 | 0.0007 |
| 372 | PIQ | IR | 11.99±9.60 | 0.109 | 0.0015 |
| | VIQ | | 12.54±9.76 | 0.051 | 0.1369 |

Table 15: Absolute IQ prediction performance by 3D-DenseNet169 in setting 1. MAE and correlation coefficient ($r$) are combined over 5-folds. For $p$-values equal to or smaller than 0.0001 are shown as 0.0001. Acronyms are used as, Exp: experiment number, I: intensity map as input, R: RAVENS map as input, and IR: combined intensity and RAVENS maps as input.

| Exp. | IQ | Data | Mean MAE | $r$ | $p$-value |
|---|---|---|---|---|---|
| 373 | | I | 11.99±9.69 | 0.060 | 0.0827 |
| 374 | PIQ | R | 11.92±9.79 | 0.080 | 0.0198 |
| 375 | | IR | 11.98±9.66 | 0.076 | 0.0267 |
| 376 | | I | 12.37±9.65 | 0.165 | 0.0001 |
| 377 | VIQ | R | 12.42±9.76 | 0.111 | 0.0012 |
| 378 | | IR | 12.38±9.90 | 0.097 | 0.0048 |
| 379 | | I | 11.70±9.06 | 0.135 | 0.0001 |
| 380 | FSIQ | R | 11.72±9.07 | 0.137 | 0.0001 |
| 381 | | IR | 11.71±9.17 | 0.092 | 0.0072 |

Table 16: Absolute IQ prediction performance by 3D-DenseNet169 in setting 2. MAE and correlation coefficient ($r$) are combined over 5-folds. For $p$-values equal to or smaller than 0.0001 are shown as 0.0001. Acronyms are used as, Exp: experiment number, I: intensity map as input, R: RAVENS map as input, and IR: combined intensity and RAVENS maps as input.

| **Exp.** | **IQ** | **Data** | **Mean MAE** | $r$ | $p$**-value** |
|---|---|---|---|---|---|
| 382 | FSIQ | I | 11.76±9.04 | 0.125 | 0.0003 |
| | PIQ | | 12.00±9.59 | 0.098 | 0.0041 |
| | VIQ | | 12.42±9.67 | 0.138 | 0.0001 |
| 383 | FSIQ | R | 11.75±9.04 | 0.123 | 0.0003 |
| | PIQ | | 11.98±9.61 | 0.113 | 0.0010 |
| | VIQ | | 12.43±9.75 | 0.119 | 0.0005 |
| 384 | FSIQ | IR | 11.74±9.08 | 0.107 | 0.0018 |
| | PIQ | | 12.02±9.63 | 0.075 | 0.0279 |
| | VIQ | | 12.43±9.83 | 0.094 | 0.0060 |

Table 17: Residual IQ prediction performance by 3D-ResNet18 in setting 1. MAE and correlation coefficient ($r$) are combined over 5-folds. For $p$-values equal to or smaller than 0.0001 are shown as 0.0001. Acronyms are used as, Exp: experiment number, I: intensity map as input, R: RAVENS map as input, and IR: combined intensity and RAVENS maps as input.

| **Exp.** | **IQ** | **Data** | **Mean MAE** | $r$ | $p$**-value** |
|---|---|---|---|---|---|
| 385 | PIQ | I | 11.37±9.58 | 0.084 | 0.0140 |
| 386 | | R | 11.40±9.53 | 0.077 | 0.0248 |
| 387 | | IR | 11.36±9.57 | 0.084 | 0.0145 |
| 388 | VIQ | I | 11.59±9.54 | 0.150 | 0.0001 |
| 389 | | R | 11.65±9.60 | 0.090 | 0.0085 |
| 390 | | IR | 11.68±9.58 | 0.081 | 0.0184 |
| 391 | FSIQ | I | 10.91±8.97 | 0.140 | 0.0001 |
| 392 | | R | 11.02±9.13 | 0.009 | 0.7829 |
| 393 | | IR | 11.01±9.04 | 0.055 | 0.1074 |

Table 18: Residual IQ prediction performance by 3D-ResNet18 in setting 2. MAE and correlation coefficient ($r$) are combined over 5-folds. For $p$-values equal to or smaller than 0.0001 are shown as 0.0001. Acronyms are used as, Exp: experiment number, I: intensity map as input, R: RAVENS map as input, and IR: combined intensity and RAVENS maps as input.

| **Exp.** | **IQ** | **Data** | **Mean MAE** | $r$ | $p$**-value** |
|---|---|---|---|---|---|

Continued on next page

**Table 18 – continued from previous page**

| Exp. | IQ | Data | Mean MAE | $r$ | $p$-value |
|---|---|---|---|---|---|
| 394 | FSIQ | I | 10.95±8.95 | 0.130 | 0.0002 |
| | PIQ | | 11.43±9.52 | 0.064 | 0.0619 |
| | VIQ | | 11.62±9.55 | 0.147 | 0.0001 |
| 395 | FSIQ | R | 11.01±8.99 | 0.058 | 0.0913 |
| | PIQ | | 11.41±9.55 | 0.058 | 0.0903 |
| | VIQ | | 11.70±9.60 | 0.057 | 0.0982 |
| 396 | FSIQ | IR | 11.06±8.93 | 0.054 | 0.1178 |
| | PIQ | | 11.43±9.56 | 0.026 | 0.4518 |
| | VIQ | | 11.69±9.56 | 0.087 | 0.0110 |

Table 19: Residual IQ prediction performance by 3D-ResNet50 in setting 1. MAE and correlation coefficient ($r$) are combined over 5-folds. For $p$-values equal to or smaller than 0.0001 are shown as 0.0001. Acronyms are used as, Exp: experiment number, I: intensity map as input, R: RAVENS map as input, and IR: combined intensity and RAVENS maps as input.

| Exp. | IQ | Data | Mean MAE | $r$ | $p$-value |
|---|---|---|---|---|---|
| 397 | PIQ | I | 11.37±9.55 | 0.087 | 0.0114 |
| 398 | | R | 11.38±9.54 | 0.081 | 0.0189 |
| 399 | | IR | 11.36±9.57 | 0.080 | 0.0199 |
| 400 | VIQ | I | 11.59±9.55 | 0.143 | 0.0001 |
| 401 | | R | 11.64±9.71 | 0.070 | 0.0413 |
| 402 | | IR | 11.69±9.62 | 0.055 | 0.1092 |
| 403 | FSIQ | I | 10.92±8.95 | 0.130 | 0.0001 |
| 404 | | R | 11.02±8.98 | 0.041 | 0.2360 |
| 405 | | IR | 11.03±8.94 | 0.097 | 0.0046 |

Table 20: Residual IQ prediction performance by 3D-ResNet50 in setting 2. MAE and correlation coefficient ($r$) are combined over 5-folds. For $p$-values equal to or smaller than 0.0001 are shown as 0.0001. Acronyms are used as, Exp: experiment number, I: intensity map as input, R: RAVENS map as input, and IR: combined intensity and RAVENS maps as input.

| Exp. | IQ | Data | Mean MAE | $r$ | $p$-value |
|---|---|---|---|---|---|
| 406 | FSIQ | I | 10.96±8.86 | 0.142 | 0.0001 |
| | PIQ | | 11.35±9.53 | 0.104 | 0.0024 |
| | VIQ | | 11.61±9.52 | 0.139 | 0.0001 |
| 407 | FSIQ | R | 11.02±8.98 | 0.060 | 0.0823 |
| | PIQ | | 11.41±9.55 | 0.083 | 0.0149 |
| | VIQ | | 11.71±9.59 | 0.060 | 0.0787 |
| 408 | FSIQ | IR | 11.05±8.94 | 0.062 | 0.0730 |
| | PIQ | | 11.42±9.55 | 0.072 | 0.0349 |
| | VIQ | | 11.70±9.61 | 0.045 | 0.1945 |

Table 21: Residual IQ prediction performance by 3D-DenseNet121 in setting 1. MAE and correlation coefficient ($r$) are combined over 5-folds. For $p$-values equal to or smaller than 0.0001 are shown as 0.0001. Acronyms are used as, Exp: experiment number, I: intensity map as input, R: RAVENS map as input, and IR: combined intensity and RAVENS maps as input.

| **Exp.** | **IQ** | **Data** | **Mean MAE** | $r$ | $p$**-value** |
|---|---|---|---|---|---|
| 409 | | I | 11.35±9.60 | 0.085 | 0.0130 |
| 410 | PIQ | R | 11.34±9.57 | 0.095 | 0.0055 |
| 411 | | IR | 11.38±9.53 | 0.094 | 0.0059 |
| 412 | | I | 11.56±9.55 | 0.152 | 0.0001 |
| 413 | VIQ | R | 11.68±9.58 | 0.102 | 0.0030 |
| 414 | | IR | 11.65±9.56 | 0.107 | 0.0017 |
| 415 | | I | 10.88±8.95 | 0.155 | 0.0001 |
| 416 | FSIQ | R | 11.02±9.02 | 0.053 | 0.1203 |
| 417 | | IR | 11.01±9.02 | 0.052 | 0.1281 |

Table 22: Residual IQ prediction performance by 3D-DenseNet121 in setting 2. MAE and correlation coefficient ($r$) are combined over 5-folds. For $p$-values equal to or smaller than 0.0001 are shown as 0.0001. Acronyms are used as, Exp: experiment number, I: intensity map as input, R: RAVENS map as input, and IR: combined intensity and RAVENS maps as input.

| **Exp.** | **IQ** | **Data** | **Mean MAE** | $r$ | $p$**-value** |
|---|---|---|---|---|---|
| | FSIQ | | 10.92±8.89 | 0.158 | 0.0001 |
| 418 | PIQ | I | 11.42±9.47 | 0.103 | 0.0027 |
| | VIQ | | 11.60±9.49 | 0.153 | 0.0001 |
| | FSIQ | | 11.04±9.01 | -0.015 | 0.6635 |
| 419 | PIQ | R | 11.42±9.55 | 0.049 | 0.1518 |
| | VIQ | | 11.70±9.65 | -0.004 | 0.9002 |
| | FSIQ | | 11.03±8.96 | 0.056 | 0.1039 |
| 420 | PIQ | IR | 11.39±9.55 | 0.086 | 0.0125 |
| | VIQ | | 11.71±9.63 | 0.009 | 0.7853 |

Table 23: Residual IQ prediction performance by 3D-DenseNet169 in setting 1. MAE and correlation coefficient ($r$) are combined over 5-folds. For $p$-values equal to or smaller than 0.0001 are shown as 0.0001. Acronyms are used as, Exp: experiment number, I: intensity map as input, R: RAVENS map as input, and IR: combined intensity and RAVENS maps as input.

| **Exp.** | **IQ** | **Data** | **Mean MAE** | $r$ | $p$**-value** |
|---|---|---|---|---|---|

Continued on next page

**Table 23 – continued from previous page**

| Exp. | IQ | Data | Mean MAE | $r$ | $p$-value |
|---|---|---|---|---|---|
| 421 | | I | 11.31±9.57 | 0.111 | 0.0012 |
| 422 | PIQ | R | 11.36±9.56 | 0.085 | 0.0131 |
| 423 | | IR | 11.37±9.53 | 0.099 | 0.0040 |
| 424 | | I | 11.54±9.50 | 0.168 | 0.0001 |
| 425 | VIQ | R | 11.66±9.59 | 0.088 | 0.0099 |
| 426 | | IR | 11.67±9.54 | 0.104 | 0.0023 |
| 427 | | I | 10.92±8.99 | 0.136 | 0.0001 |
| 428 | FSIQ | R | 11.01±8.96 | 0.089 | 0.0098 |
| 429 | | IR | 11.00±9.01 | 0.046 | 0.1783 |

Table 24: Residual IQ prediction performance by 3D-DenseNet169 in setting 2. MAE and correlation coefficient ($r$) are combined over 5-folds. For $p$-values equal to or smaller than 0.0001 are shown as 0.0001. Acronyms are used as, Exp: experiment number, I: intensity map as input, R: RAVENS map as input, and IR: combined intensity and RAVENS maps as input.

| Exp. | IQ | Data | Mean MAE | $r$ | $p$-value |
|---|---|---|---|---|---|
| | FSIQ | | 10.96±8.88 | 0.142 | 0.0001 |
| 430 | PIQ | I | 11.38±9.50 | 0.110 | 0.0013 |
| | VIQ | | 11.63±9.50 | 0.142 | 0.0001 |
| | FSIQ | | 11.02±8.99 | 0.041 | 0.2278 |
| 431 | PIQ | R | 11.40±9.56 | 0.062 | 0.0701 |
| | VIQ | | 11.71±9.67 | 0.019 | 0.5716 |
| | FSIQ | | 11.00±8.95 | 0.088 | 0.0103 |
| 432 | PIQ | IR | 11.38±9.54 | 0.092 | 0.0074 |
| | VIQ | | 11.72±9.61 | 0.041 | 0.2316 |
